# Neuronal UBE3A loss engages a TNF-driven neuron-to-microglia axis that promotes synapse remodeling

**DOI:** 10.64898/2026.05.27.728269

**Authors:** Xin Yang, Xu Zhang, Yu-Wen Alvin Huang

## Abstract

Neuron–microglia communication shapes neuroimmune homeostasis and circuit maturation, yet the neuron-derived cues that tune microglial inflammatory state and phagocytic programs remain incompletely defined. The E3 ubiquitin ligase UBE3A is critical for brain development and synaptic function, and altered UBE3A dosage or activity is linked to neurodevelopmental phenotypes with emerging connections to inflammation. Using Angelman syndrome (AS), where UBE3A loss is largely neuron-selective due to imprinting, as a tractable framework, we investigated how neuronal UBE3A deficiency engages microglial programs.

In human iPSC-derived neuron–microglia co-cultures, neuronal UBE3A depletion induced a TNF-associated secretory signature and was accompanied by increased microglial inflammatory markers, including complement components. Neuron-conditioned media was sufficient to elicit microglial cytokine/complement transcripts, supporting a role for soluble neuron-derived cues. In these co-cultures, neuronal UBE3A loss increased synapse engulfment by microglia, which was reduced by TNF receptor pathway inhibition with the TNFR1 antagonist R7050 alongside decreased microglial inflammatory readouts. In vivo, juvenile AS mouse hippocampus showed early microglial dysregulation, and reanalysis of a maternal-deletion AS pig single-nucleus RNA-seq dataset revealed enrichment of microglial activation and phagosome/lysosome pathways. Together, these findings implicate a TNF-linked neuron-to-microglia signaling axis downstream of neuronal UBE3A loss that can be pharmacologically modulated during development.

## Introduction

Precise brain wiring depends on continuous crosstalk between neurons and microglia, the resident macrophages of the central nervous system. During early postnatal development, microglia integrate neuron-derived “find-me/eat-me” cues, activity-dependent signals, and local inflammatory mediators to survey circuits and remove unnecessary synapses, thereby refining connectivity in a manner essential for normal cognition (*1–3*). Landmark studies have shown that microglia physically engulf synaptic material during developmental pruning and that complement-dependent pathways (e.g., C3-CR3 signaling) help tune activity-dependent circuit refinement (*4–6*); disruption of these processes has been implicated in schizophrenia (*7*) and autism spectrum disorders (*8–10*). Yet, despite rapidly expanding appreciation of microglia as sculptors of neural circuits and drivers of brain disease risk, the upstream neuronal programs that restrain microglial activation state and calibrate synaptic engulfment during development remain incompletely defined.

Angelman syndrome (AS) provides a uniquely powerful genetic framework to dissect neuron-to-microglia signaling because it features a striking, cell-type-biased disruption of the same gene across the developing brain. AS is caused by loss of the maternally inherited *UBE3A* allele in neurons, owing to neuron-specific genomic imprinting in which the paternal allele is transcriptionally silenced by a long antisense transcript (UBE3A-ATS); in contrast, glial populations largely retain biallelic UBE3A expression (*11–13*). This genetic architecture creates an experimental “separation of variables”: neuronal UBE3A loss can be interrogated as an upstream driver while microglia remain comparatively intact at the locus, enabling focused tests of neuron-derived signals that instruct microglial states. Mechanistically, UBE3A is a HECT-family E3 ligase with established roles in synaptic plasticity and learning-related circuit function, and AS brains show synaptic and spine abnormalities that have historically been interpreted primarily through neuron-intrinsic mechanisms (*14–16*). Notably, aberrant UBE3A dosage and signaling are implicated in a broader spectrum of brain disorders beyond AS, including autism-associated 15q11–q13 duplication/UBE3A overexpression syndromes (*17, 18*) and emerging links to neurodegenerative and neuroinflammatory pathways, highlighting UBE3A as a convergent regulator of circuit stability across the lifespan (*19*). Whether, and how, neuronal UBE3A deficiency disrupts neuron-to-microglia communication to contribute to synaptic pathology remains largely unresolved.

A major conceptual gap concerns pro-inflammatory cytokine signaling originating from neurons. Microglia–astrocyte inflammatory crosstalk has been extensively characterized, including pathways in which activated microglia release cytokines that reprogram astrocytes into reactive states with downstream effects on neuronal survival and synaptic function (*20, 21*). In contrast, the conditions under which neurons themselves initiate cytokine-like programs that meaningfully tune microglial activation and phagocytic behavior - particularly during neurodevelopment - are less well understood. Tumor necrosis factor-α (TNF-α) is classically associated with immune and glial signaling in the CNS, and is a potent modulator of synaptic physiology and inflammatory state across cell types (*22*). Neuronal TNF-α expression has also been reported in several stress and injury contexts (*23–26*), raising the possibility that aberrant neuronal TNF-α induction could function as an intercellular cue that biases microglial state and synapse remodeling.

Here, we use a reduced human iPSC-derived neuron-microglia platform (*27–29*) together with *in vivo* analyses in juvenile AS mouse hippocampus (*30, 31*) and reanalysis of a maternal-deletion AS pig brain single-nucleus RNA-seq dataset (*32, 33*) to examine how neuronal UBE3A deficiency engages microglial inflammatory and phagocytic programs. We show that neuronal UBE3A knockdown induces a TNF-associated inflammatory signature and that neuron-conditioned media is sufficient to elicit microglial cytokine and complement transcripts. In co-culture, neuronal UBE3A deficiency is accompanied by increased microglia-associated presynaptic marker signal, a proxy for synaptic material associated with microglia, alongside microglial inflammatory readouts; these phenotypes are reduced by pharmacologic inhibition of TNF receptor signaling. In vivo, juvenile AS mouse hippocampus exhibits early alterations in microglial abundance and activation-associated markers, and pig snRNA-seq microglial clusters show enrichment of phagosome/lysosome and antigen-processing programs consistent with elevated phagocytic states. Together, our findings implicate a TNF-linked neuron-to-microglia signaling axis downstream of neuronal UBE3A loss and motivate further work to define the causal chain from neuronal cytokine programs to microglial synapse remodeling during development.

## RESULTS

### Neuronal UBE3A deficiency induces a TNF-associated neuronal inflammatory signature and is accompanied by microglial inflammatory activation

To dissect neuron–microglia interactions downstream of neuronal UBE3A loss, we leveraged our established human iPSC-based neuroinflammation platform (*27–29*), consisting of induced neurons (iNs) and iPSC-derived microglia-like cells (iMG) generated from the common reference line KOLF2.J1 (*34*). This reduced neuron-microglia circuit enabled us to test whether neuronal UBE3A depletion engages a neuron-intrinsic inflammatory program and whether neuron-derived cues are sufficient to shift microglial inflammatory state. To model the neuron-biased UBE3A deficiency observed in Angelman syndrome (AS), we performed RNAi-mediated knockdown of UBE3A in iNs (siUBE3A) compared to a non-targeting control (siNC).

We first asked whether acute UBE3A depletion produces broad cell-autonomous transcriptional changes in neurons. Using qPCR, we assessed pro-inflammatory cytokines (C3, IL1B, TNFA), synaptic and neuronal identity genes (SYN1, SYT1, GABRA1, GRIA1), activity-dependent genes (ARC, CREB1, cFOS), and genes related to extracellular remodeling and cytokine processing, including PTX3 (an inflammatory pentraxin implicated in synapse remodeling), ADAM10 (a sheddase that can process multiple cell-surface substrates, including pro-TNF family ligands), and MMP9 (a matrix metalloprotease linked to activity-and inflammation-associated remodeling). Notably, at this timepoint, neuronal UBE3A knockdown did not broadly suppress synaptic or activity-dependent transcripts, suggesting an early selective transcriptional shift rather than generalized downregulation of synaptic machinery (**Fig. 1A**). In contrast, we observed a marked induction of TNFA mRNA (and a more modest increase in MMP9), consistent with engagement of a neuron-associated inflammatory program (**Fig. 1A**). Because many UBE3A-linked synaptic phenotypes arise through post-transcriptional proteostasis and activity-dependent remodeling, broad synaptic mRNA changes may be modest at early timepoints, and transcript levels may not reflect synaptic function directly. We note that functional synaptic impairment was not directly assessed. Because many UBE3A-linked synaptic phenotypes arise through post-transcriptional proteostasis and activity-dependent remodeling, broad synaptic mRNA changes may be modest at early timepoints, and transcript levels may not reflect synaptic function directly..

**Figure 1.**
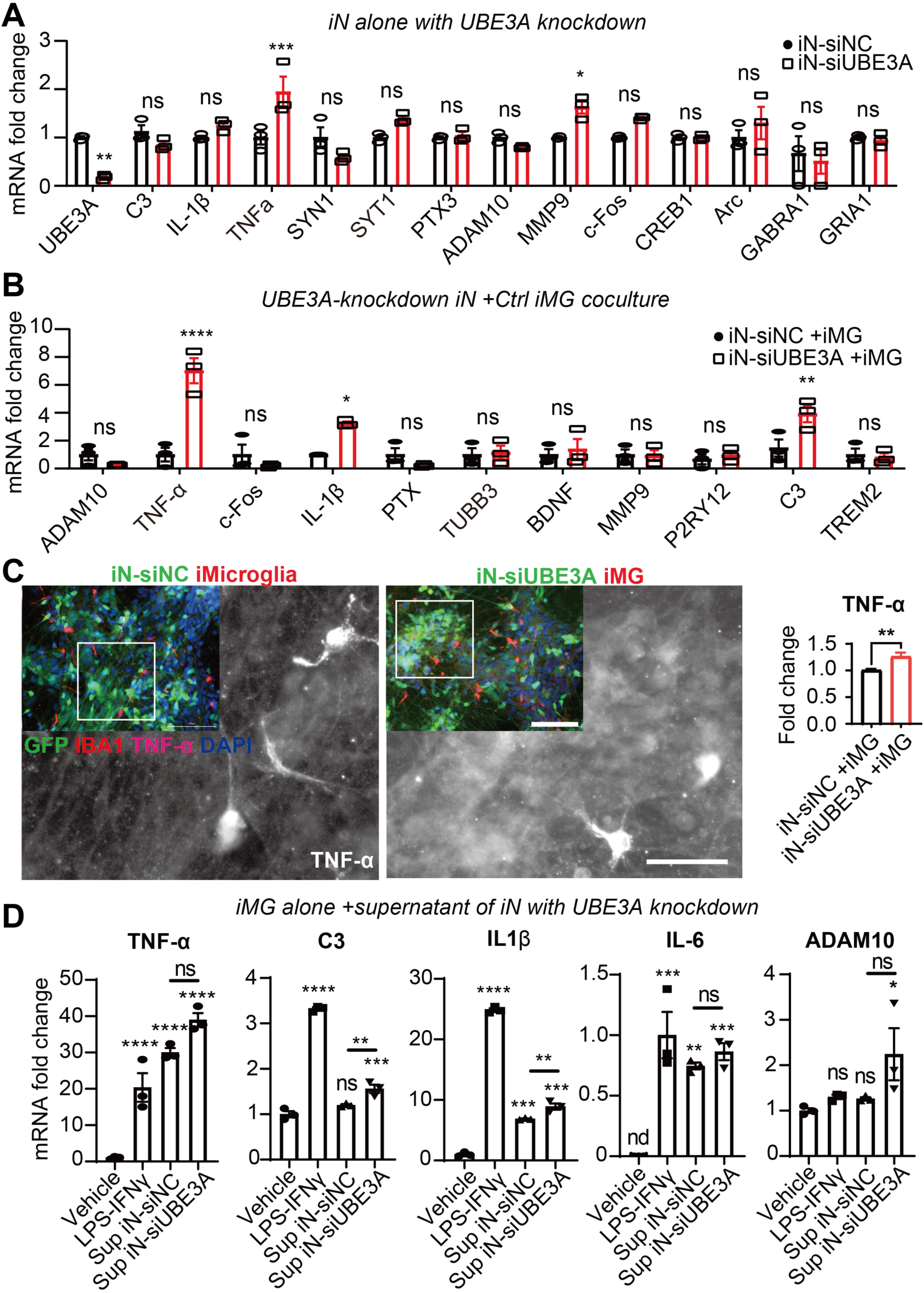
Neuronal UBE3A depletion induces TNF-α and primes microglia through soluble neuron-derived signals. (A) qPCR analysis of mature human induced neurons (iN) derived from the KOLF2 (KOLF2.J1) reference iPSC line and transduced with non-targeting control siRNA (siNC) or siRNA targeting UBE3A (siUBE3A). Transcripts assessed include pro-inflammatory mediators (C3, TNFα and IL-1β), synaptic/neuronal genes (SYN1, SYT1, PTX3, GABRA1, GRIA1), protease/sheddase and remodeling-associated genes (ADAM10, MMP9), and activity-dependent genes (ARC, CREB1, cFOS). (B) qPCR analysis of iMG co-cultured with iN-siNC or iN-siUBE3A, assessing pro-inflammatory genes (TNFα, IL-1β and C3), microglial marker genes (P2RY12, TREM2), neuronal marker (TUBB3), and additional candidate signaling/remodeling genes (ADAM10, PTX3, MMP9, BDNF). (C) Representative confocal images and quantification of TNF-α immunoreactivity in iN–iMG co-cultures. Neurons express GFP from the FUGW lentivirus (green) and microglia are labeled with IBA1 (red); TNF-α is shown in magenta and DAPI in blue. (D) qPCR analysis of iMG treated with conditioned media from iN-siNC or iN-siUBE3A cultures (Sup iN-siNC and Sup iN-siUBE3A, respectively) or with LPS+IFN-γ as a positive control, measuring induction of TNFα, C3, IL1B, IL6, and ADAM10.

Because TNF-α is commonly viewed as a glia-associated cytokine, we next contextualized our neuronal TNFA induction by surveying published cell type–resolved brain transcriptomic datasets. We surveyed published cell-type–resolved transcriptomic datasets and found that Tnf/TNF expression is highest in microglia in both mouse (*35*) and human (*36*) brains, while remaining detectable in neurons (**Fig. S1A-B**). These data support TNF-α as a canonical microglial cytokine while also underscoring the plausibility that neuronal TNF-α transcripts can contribute to local signaling under specific contexts..

We then asked whether neuronal UBE3A depletion is accompanied by changes in microglial inflammatory state in co-culture. iMG were co-cultured with iN-siNC or iN-siUBE3A, and qPCR was used to measure inflammatory readouts (TNFA, IL1B, C3), microglial markers (P2RY12, TREM2), and additional modulators of synaptic and trophic support (ADAM10, PTX3, MMP9, BDNF) alongside the neuronal marker TUBB3. Co-cultures containing UBE3A-deficient neurons showed increased expression of inflammatory markers, including TNFA and C3 (**Fig. 1B**), consistent with a shift toward a more inflammatory microglial state in the presence of UBE3A-deficient neurons. To complement these transcript-level observations, we performed confocal immunofluorescence imaging of co-cultures labeled for neurons (GFP), microglia (IBA1), and TNF-α. TNF-α immunoreactivity was increased in co-cultures containing UBE3A-deficient neurons (**Fig. 1C**).

To determine whether microglial inflammatory induction requires direct cell–cell contact or can be elicited by soluble neuron-derived cues, we treated iMG with neuronal conditioned media (supernatant) from control or UBE3A-deficient neurons. As a positive control for microglial responsiveness, LPS + IFNγ robustly induced TNFA, C3, IL1B, and IL6, confirming immune competency of the iMG (Fig. 1D). Neuronal supernatant from control iNs (Sup iN-siNC) induced measurable microglial cytokine transcripts (notably IL1B and IL6, with TNFA induction depending on the readout), consistent with the concept that neuron-derived soluble factors can influence microglial inflammatory tone under baseline conditions. In comparison, supernatant from UBE3A-deficient neurons (Sup iN-siUBE3A) produced moderate but statistically significant increases in select inflammatory transcripts relative to control supernatant (**Fig. 1D**). We therefore interpret these results as evidence that soluble neuron-derived cues contribute to microglial activation state, with neuronal UBE3A loss modestly shifting the secretome in a direction associated with increased microglial inflammatory readouts. To validate the conditioned-media effect in an orthogonal microglial model, we repeated the supernatant stimulation paradigm using HMC3 cells, an immortalized human microglial line widely used for mechanistic perturbation and reproducible inflammatory assays (*37*). Consistent with the iMG results, UBE3A-deficient neuronal supernatant elicited stronger induction of inflammation-associated transcripts in HMC3 cells compared with control neuronal supernatant (**Fig. S2**), further supporting a role for soluble neuron-derived signals.

Because UBE3A is broadly expressed across brain cell types, yet its cell-autonomous role in microglia remains poorly defined, we next asked whether altering UBE3A levels within microglia can directly influence microglial identity and inflammatory state. Although AS is primarily driven by neuron-selective loss of UBE3A (with glial UBE3A largely preserved), we performed siRNA-mediated UBE3A knockdown in human iPSC-derived microglia-like cells (iMG) and in the HMC3 microglial cell line to probe intrinsic microglial effects. UBE3A depletion did not measurably disrupt expression of core microglial markers (e.g., IBA1 and TREM2) or basal lipid-droplet burden, but it increased pro-inflammatory cytokine transcripts (TNFα and IL-1β) in iMG and sensitized HMC3 cells to exaggerated inflammatory induction upon LPS stimulation (**Fig. S3**). These results suggest that microglial UBE3A contributes to restraining inflammatory tone and reactivity, providing a mechanistic context for how neuron-derived signals may further amplify microglial activation in the setting of neuronal UBE3A deficiency.

Together, these experiments implicate a TNF-associated neuron-to-microglia signaling cascade downstream of neuronal UBE3A loss. In this reduced human platform, neuronal UBE3A depletion did not broadly suppress synaptic or activity-dependent neuronal transcripts at the tested timepoint, but instead induced a selective inflammatory signature dominated by TNFA. Co-culture and conditioned-media paradigms indicate that neuron-derived cues are sufficient to engage microglial inflammatory readouts, and an orthogonal microglial model (HMC3) supports the robustness of supernatant-driven activation. These findings motivate subsequent analyses of how TNF-linked microglial activation states relate to synapse-associated phenotypes in this system.

Data are mean ± SEM. n = 3–5 independent experiments. Statistics: (A,B) two-way ANOVA; (C) two-tailed unpaired t test; (D) one-way ANOVA; selected post-hoc comparisons shown here, vs. control conditions (A, iN-siNC; B and C, iN-siNC +iMG; D, Vehicle) and in (D), Sup iN-siNC vs. Sup iN-siUBE3A. * p< 0.05, ** p< 0.01, *** p< 0.001, **** p< 0.0001, ns, not significant; nd, not detected.

### *In vivo* Validation of microglial activation in rodent and porcine models of Angelman syndrome

Our reduced human co-culture system identified neuron-derived inflammatory cues, dominated by TNF-α, as a sufficient trigger for microglial priming. We next asked whether comparable microglial dysregulation emerges in vivo during early postnatal development, when microglia actively sculpt hippocampal circuits through synapse remodeling. AS is particularly well-suited for this question because neuronal UBE3A is selectively lost (via neuron-specific imprinting), enabling a focused test of how a neuron-intrinsic genetic lesion can secondarily reshape microglial state. We therefore examined hippocampal tissue from the AS transgenic mice (*30, 31*) at the early juvenile stage (P14) aligned with robust circuit refinement.

Immunohistological analysis of juvenile hippocampus confirmed UBE3A signal is reduced prominently in neuronal layers in AS tissue (**Fig. 2A**), establishing the *in vivo* substrate for neuron-driven inflammatory signaling. We observed a broader inflammatory milieu within AS hippocampus: immunostaining revealed increased TNF-α and C3 signals overall, and importantly, these increases were even more pronounced when analyses were restricted to SYN1+ neuronal/synaptic regions (**Fig. S4**). Surprisingly, quantification of IBA1+ microglia revealed a decrease in microglial density in the developing AS hippocampus (**Fig. 2B**), suggesting that neuronal UBE3A loss perturbs microglial abundance and/or maintenance during this developmental window. Despite fewer microglia, remaining AS microglia exhibited elevated CD68 signal within IBA1+ cells (**Fig. 2C**). Because CD68 is a lysosomal/phagolysosomal protein widely used as a readout of phagocytic/activated microglial states, this pattern is consistent with a shift toward heightened microglial activation *in vivo* (*38*).

**Fig. 2.**
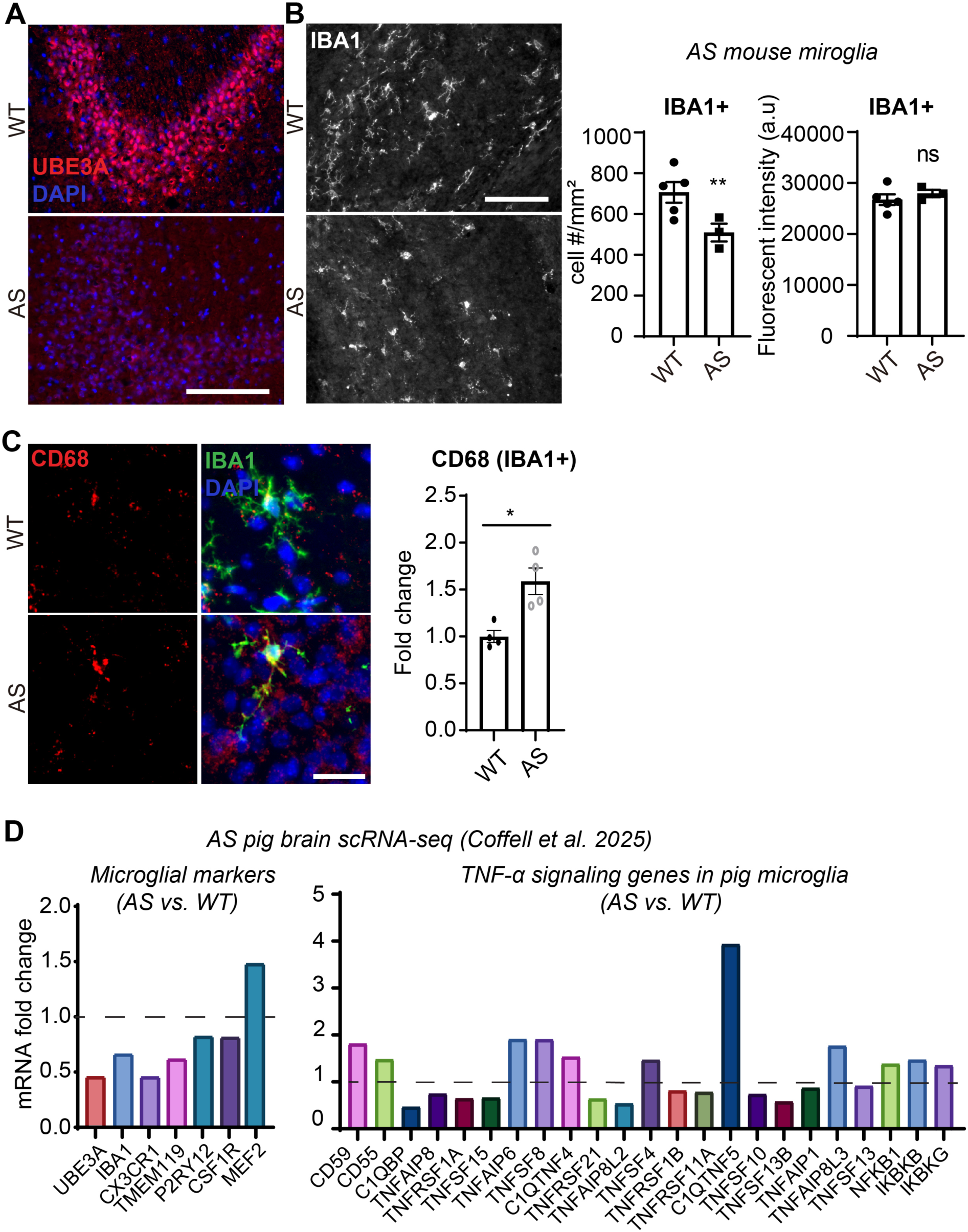
I*n vivo* validation of microglial dysregulation and activation in Angelman syndrome mouse and pig models. (A) Representative immunofluorescence images of hippocampal sections from P14 WT and AS mice stained for UBE3A (red) and nuclei (DAPI, blue), illustrating reduced UBE3A immunoreactivity most prominently in neuronal layers in AS. Scale bar, 100 μm. (B) Representative images of hippocampal IBA1 immunostaining (microglia) in WT and AS mice, with quantification of microglial density and IBA1 signal intensity. n = 5 WT and 3 AS animals. Statistics: two-tailed unpaired t test. ** p< 0.01. Scale bar, 200 μm. (C) Representative higher-magnification images of IBA1 (microglia) and CD68 (lysosomal/phagolysosomal marker enriched in activated/phagocytic microglia) staining in WT and AS hippocampus, with quantification of microglial CD68 signal (CD68 within IBA1⁺ microglia). n = 5 animals per group. Statistics: two-tailed unpaired t test. * p< 0.05. Scale bar, 50 μm. (D) Reanalysis of a published single-nucleus RNA-seq dataset from a maternal-deletion AS pig model at P10. Microglial nuclei were subsetted and compared between AS and WT. Left, differential expression of microglial identity/homeostatic markers (including TMEM119 and P2RY12) and UBE3A. Right, relative induction of TNF-α–responsive signaling genes in AS microglia compared with WT, consistent with an activated microglial state.

To test whether this microglial signature generalizes beyond rodents, and also to leverage a brain architecture closer to humans, we reanalyzed a published maternal-deletion AS pig single-nucleus RNA-seq dataset (Coffell *et al.*, 2025)(*32*) (**Fig. 2D**). Large-animal AS pig models have been developed to better mirror early developmental features and neurological trajectory that can be challenging to capture in rodents (*33*). In the young pig brain snRNA-seq dataset, UBE3A expression in microglia was reduced to approximately half of WT (**Fig. 2D**, left), consistent with retained paternal-allele expression outside neurons, and microglial homeostatic programs (including TMEM119/P2RY12-class markers) were broadly downshifted alongside induction of TNF-associated signaling genes (**Fig. 2D**). Moreover, when we subsetted the same dataset to excitatory neurons, differential expression followed by KEGG enrichment analysis revealed significant activation of the TNF-α signaling pathway (**Fig. S5A**), and multiple TNF-α pathway–associated genes were significantly upregulated in AS excitatory neurons compared with WT (**Fig. S5B**). These findings reinforce the conclusion that TNF-α–linked inflammatory programs are engaged not only in microglia but also within the neuronal compartment *in vivo*.

Collectively, these cross-platform and cross-species data indicate that neuronal UBE3A loss is coupled to microglial dysregulation early in development, with evidence for an activated/lysosome-enriched microglial state emerging by P14. This early timing supports a model in which aberrant microglial activation could contribute to initial circuit vulnerabilities - potentially by biasing microglial synapse remodeling toward excessive elimination - motivating our next section directly testing whether microglia show altered engulfment of synaptic material in AS.

Data are presented as mean ± SEM.

### Neuronal UBE3A loss is associated with phagocytic microglial programs and increased microglia-associated synaptic material that is modulated by TNF receptor inhibition

Having established that neuronal UBE3A deficiency engages a TNF-associated neuronal inflammatory signature and is accompanied by microglial inflammatory induction, we next asked whether this signaling context maps onto a core microglial effector program relevant to circuit maturation. Microglia regulate synapse elimination during development through tightly controlled phagocytic and lysosomal pathways (*1–3*), and multiple neurodevelopmental disease models implicate altered microglial pruning programs in circuit dysfunction, particularly in the context of autism (*8–10*). We therefore sought evidence for coordinated induction of phagocytic/lysosomal gene modules in vivo and then tested, in our reduced human co-culture platform, whether neuronal UBE3A loss alters the association of synaptic markers with microglia - a commonly used proxy for microglia-associated synaptic material.

To strengthen the *in vivo* relevance and to leverage a large-animal dataset closer to human brain organization, we continued our reanalysis of the maternal-deletion AS pig snRNA-seq dataset (*32*) by focusing specifically on the microglial cluster. Pathway enrichment analysis of differentially expressed genes (AS vs. WT) revealed significant upregulation of gene sets involved in phagosome and lysosome function, antigen processing and presentation, and broader immune-response modules (**Fig. 3A**), consistent with microglia engaging a cargo-processing and phagocytic program. Examination of individual genes further highlighted broad dysregulation of phagocytosis-and lysosome-associated transcripts in AS pig microglia (**Fig. 3B**). A complementary KEGG gene–pathway network view emphasized that these enriched terms are driven by coordinated changes in complement components (e.g., C1QA/B/C), Fc receptor genes (FCGR1A/FCGR3A), innate sensors (CD14, LY96), and lysosomal/phagolysosomal machinery (CD63/CD68, ATP6V0/ATP6V1 subunits, CTSS/CTSV), alongside swine leukocyte antigen (SLA) class I genes as indicators of antigen presentation (**Fig. S6A**). In parallel, comparison to an independent RNA-seq dataset from TNF-α–stimulated *Ube3a*−/− versus *Ube3a*+/+ mouse embryonic fibroblasts (*39*) showed enrichment of canonical TNF/NF-κB and cytokine-chemokine signaling modules (**Fig. S6B-C**), supporting the broader linkage between UBE3A loss and TNF-responsive inflammatory gene networks.

**Fig 3.**
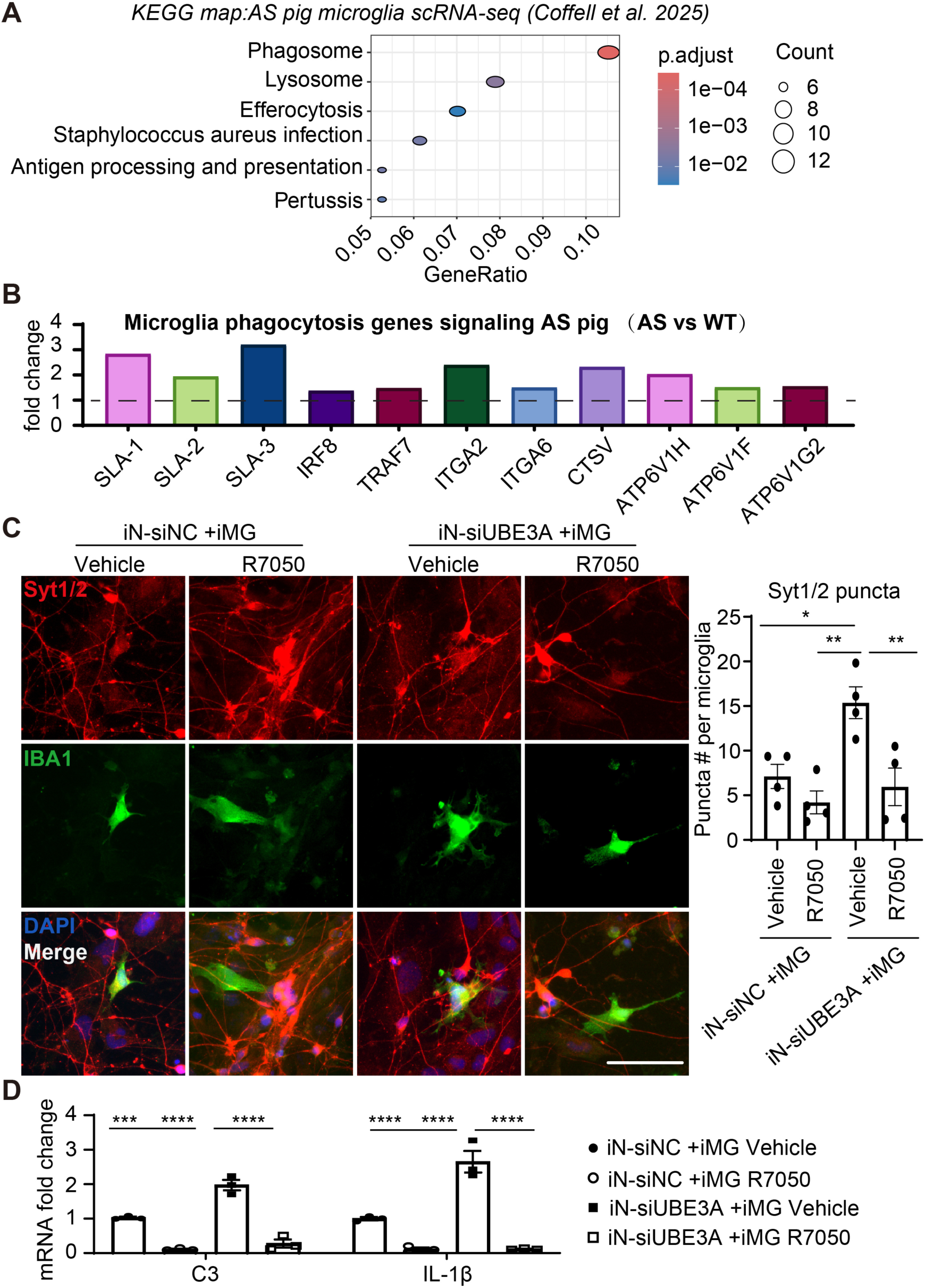
Neuronal UBE3A loss promotes a phagocytic microglial program and hyperactive synaptic engulfment via TNFα signaling. (A) KEGG pathway enrichment analysis of differentially expressed genes (AS vs. WT) in microglia extracted from a published maternal-deletion AS pig brain single-nucleus RNA-seq dataset (Coffell et al., 2025). Enriched pathways include phagosome and lysosome programs, antigen processing and presentation, and infection/immune-response pathways, consistent with an activated, phagocytic microglial state. (B) Bar-graph depiction of significantly altered microglial genes involved in phagocytosis/engulfment and lysosomal processing in AS pig microglia compared with WT (Coffell *et al.*, 2025). (C) Representative confocal images of human iN–iMG co-cultures under four conditions: iN-siNC + iMG treated with vehicle or R7050, and iN-siUBE3A + iMG treated with vehicle or R7050. Neurons were immunostained for the presynaptic marker synaptotagmin-1/2 (Syt1/2), and microglia were labeled with IBA1. Right, quantification of Syt1/2 puncta localized within IBA1+ microglia as a proxy for microglial synaptic engulfment. Neuronal UBE3A knockdown increased synaptic material within microglia, while R7050 selectively reduced over-engulfment in iN-siUBE3A co-cultures without significantly altering baseline engulfment in control co-cultures. n = 4 independent cultures. Scale bar, 50 μm. (D) qPCR analysis of inflammatory/microglial activation markers (C3 and IL1B) in the same four co-culture conditions as in (C). R7050 reduced C3 and IL1B expression, consistent with suppression of TNFα-driven microglial activation downstream of neuronal UBE3A loss. n = 3 independent cultures.

We then tested, in a controlled human iN-iMG system, whether neuronal UBE3A deficiency is accompanied by increased microglia-associated synaptic marker signal. iMG were co-cultured with iN-siNC or iN-siUBE3A and immunostained for IBA1 and the presynaptic marker synaptotagmin-1/2 (Syt1/2). We quantified the number of Syt1/2 puncta overlapping with IBA1+ microglia as a proxy for microglia-associated synaptic material (**Fig. 3C**). Importantly, this 2D colocalization metric cannot distinguish true internalization from close apposition or surface association and is therefore interpreted as an association readout rather than definitive engulfment; this limitation is discussed further below. Under these conditions, neuronal UBE3A knockdown was accompanied by increased Syt1/2 puncta overlapping with microglia compared with control co-cultures (**Fig. 3C**). To test whether this excessive engulfment is driven by TNF signaling, we used R7050, a small-molecule antagonist that inhibits TNF receptor signaling by disrupting TNFR1 adaptor interactions and downstream signaling cascades (*40*). R7050 had minimal effect on baseline synaptic marker association in control co-cultures, but it reduced the elevated Syt1/2–microglia overlap observed in iN-siUBE3A co-cultures (**Fig. 3C**), suggesting that TNF receptor pathway activity contributes to the microglia-associated synaptic marker phenotype downstream of neuronal UBE3A loss. In parallel, qPCR analysis showed that R7050 reduced expression of microglial inflammatory readouts C3 and IL1B in co-cultures (**Fig. 3D**), cconsistent with TNF receptor signaling functioning as a major modulator of microglial inflammatory state in this system.

Together, these cross-species and human co-culture data support a model in which neuronal UBE3A loss engages TNF-linked inflammatory signaling and is associated with microglial transcriptional programs enriched for phagosome/lysosome pathways, alongside increased microglia-associated synaptic marker signal in vitro. While additional orthogonal approaches would be needed to further corroborate synaptic internalization, our data support TNF/TNFR signaling as a tractable control point that dampens microglial inflammatory activation and reduces microglia-associated synaptic marker phenotypes downstream of neuronal UBE3A deficiency.

Data are mean ± SEM. Statistics, two-way ANOVA for (C, D).

### TNF receptor pathway inhibition dampens neuron-conditioned microglial activation and normalizes inflammatory outputs

Our preceding data indicate that neuronal UBE3A loss induces a TNF-α–centered secretory program that is sufficient to prime microglia and drive excessive synaptic engulfment. We next asked whether pharmacologic blockade of TNF receptor signaling can broadly normalize microglial inflammatory activation driven by neuron-derived cues in our reduced human system. To focus on the non–cell-autonomous component, we treated microglia with conditioned media from control or UBE3A-deficient neurons (Sup iN-siNC vs. Sup iN-siUBE3A) in the presence of vehicle or the TNF pathway inhibitor R7050, and quantified inflammatory outputs at the protein level.

Immunoblotting of iPSC-derived microglia-like cells (iMG) exposed to neuronal supernatant showed that conditioned media from UBE3A-deficient neurons increased TNF-α and C3 protein, including multiple C3 immunoreactive species, compared with control neuronal supernatant (**Fig. 4A**). R7050 suppressed these elevations, bringing TNF-α and C3 signals toward baseline without altering microglial UBE3A levels, consistent with an action downstream of TNF receptor signaling rather than on UBE3A expression itself (**Fig. 4A**). These results complement our transcript-level readouts and support the conclusion that neuron-derived factors produced upon neuronal UBE3A depletion engage a TNF-dependent amplification loop in microglia that drives inflammatory protein outputs.

**Fig 4.**
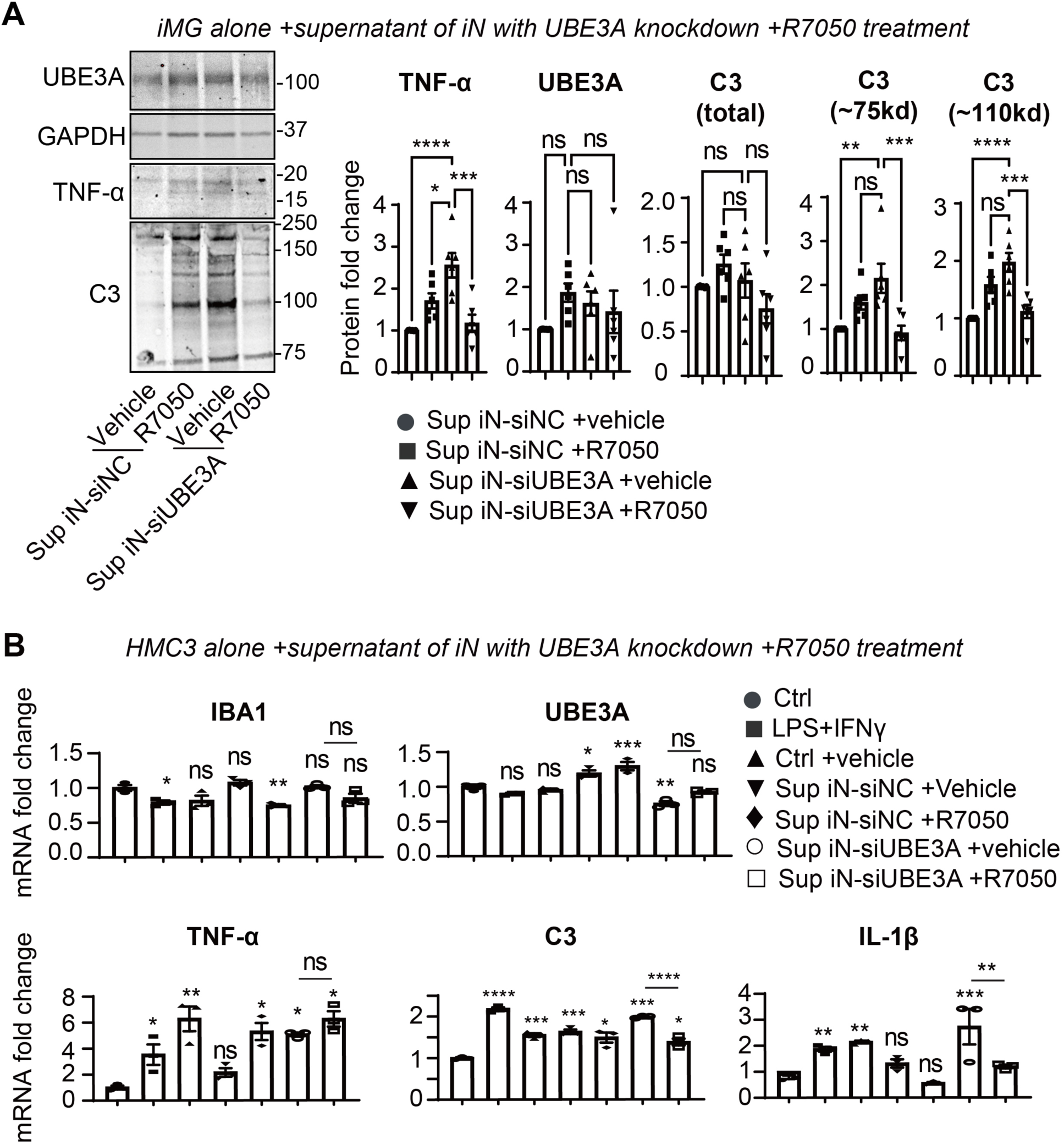
Pharmacological inhibition of TNF-α signaling normalizes the microglial inflammatory response to the UBE3A-deficient neuronal secretome. (A) Representative immunoblots and densitometric quantification of protein expression in human iMG treated with neuronal conditioned media (supernatant) from control neurons (Sup iN-siNC) or UBE3A-deficient neurons (Sup iN-siUBE3A), in the presence of vehicle or the TNF receptor signaling antagonist R7050. Blots were probed for UBE3A, TNF-α, and C3, with GAPDH as a loading control. For quantification, band intensities were first normalized to GAPDH for each lane and then expressed relative to the Sup iN-siNC + vehicle condition (set to 1.0). Quantification is shown for TNF-α, UBE3A, and C3 (total signal) as well as two prominent C3 immunoreactive species (∼75 kDa and ∼110 kDa), indicating that neuronal UBE3A deficiency elevates inflammatory protein outputs in iMG and that R7050 suppresses this response. n = 4 independent experiments. (B) qPCR analysis of the HMC3 human microglial cell line treated with: (i) control media (Ctrl), (ii) LPS+IFNã (1000/20 ng/mL, 6 h) as a positive control for microglial activation, or (iii) neuronal conditioned media from Sup iN-siNC or Sup iN-siUBE3A ± R7050 (24 h). Transcripts analyzed include microglial marker/identity genes (IBA1, UBE3A) and inflammatory readouts (TNFA, C3, IL1B). Neuron-conditioned media induced inflammatory gene expression, which was further enhanced by UBE3A-deficient neuronal supernatant; R7050 attenuated this induction, consistent with TNF-dependent control of microglial inflammatory activation in response to neuron-derived cues. n = 3 independent experiments. Data are presented as mean ± SEM. Statistics: one-way ANOVA; selected post-hoc comparisons shown here, in (D) vs. Ctrl condition and Sup iN-siUBE3A +vehicle vs. +R7050. * p< 0.05, ** p< 0.01, *** p< 0.001, **** p< 0.0001; ns, not significant.

We then tested whether this pattern generalizes to an orthogonal microglial model and whether R7050 can dampen canonical inflammatory transcriptional programs induced by neuron-derived cues. In the iMG, LPS+IFNγ produced the expected induction of inflammatory transcripts, confirming responsiveness.

Exposure to neuronal conditioned media induced TNFα, C3, and IL-1β, with stronger induction triggered by supernatant from UBE3A-deficient neurons, and R7050 attenuated these inflammatory transcripts (**Fig. 4B**). Notably, suppression was observed not only in the UBE3A-deficient neuron supernatant condition but also in control neuron supernatant conditions, consistent with TNF signaling functioning as a major integrator of neuron-derived cues that shape microglial inflammatory tone in this system (**Fig. 4B**).

Finally, while our central model emphasizes neuron-initiated signaling, we asked whether R7050 can also mitigate inflammatory protein induction caused by cell-autonomous UBE3A depletion in microglia. In iMG transduced with siUBE3A, R7050 reduced TNF-α and C3 protein readouts without restoring UBE3A protein (**Fig. S7**), suggesting that TNF receptor signaling contributes to inflammatory amplification downstream of microglial intrinsic perturbation as well. Importantly, these effects were modest compared with the strong normalization observed when microglia were driven by conditioned media from UBE3A-deficient neurons, reinforcing that the dominant driver in our model is the non–cell-autonomous neuron-to-microglia inflammatory axis.

Together, these data show that TNF receptor pathway inhibition is sufficient to dampen microglial inflammatory activation elicited by secreted factors from UBE3A-deficient neurons and to reduce downstream complement/cytokine outputs. This positions neuronal TNF-α/TNFR signaling as a therapeutically tractable control point to rebalance microglial inflammatory state and, by extension, microglial synaptic remodeling programs downstream of neuronal UBE3A loss.

Data are mean ± SEM; ns, not significant. Statistics, one-way ANOVA; * p<0.05, ** p<0.01.

## Discussion

Our study addresses a central question in neurodevelopmental biology: how do neuron-intrinsic perturbations instruct microglial state and synaptic remodeling during circuit assembly? Using complementary human iPSC-derived neuron–microglia systems and *in vivo* validation in AS mouse and pig models, we identify a neuron-initiated inflammatory axis in which neuronal UBE3A deficiency induces a TNF-α–centered secretory program that primes microglia, elevates microglial inflammatory outputs (including complement and cytokines), and promotes excessive synaptic engulfment. These findings connect neuron-to-microglia communication to a concrete developmental effector function - synapse elimination - building on foundational work showing that microglia physically prune synapses during development (*1, 4*) and that dysregulated pruning is implicated in neurodevelopmental disease (*8–10*). Mechanistically, our data position TNF signaling as a major integrator of neuron-derived cues that shifts microglia toward a phagocytic/lysosome-enriched state, consistent with broader frameworks of neuron-glia signaling in synapse elimination and complement-associated synapse loss.

Neuroinflammation is often conceptualized as a glia-dominant process, with microglia and astrocytes serving as principal sources of cytokines that modulate neuronal excitability, synaptic strength, and memory (*41–43*). However, accumulating evidence indicates that neurons can actively participate in inflammatory signaling, including expression of TNF-α and other cytokines in response to injury or stress, with functional consequences for neural outcomes (*23–26*). For example, cortical neurons were found to express TNF-α in the brain before immune cells under ischemia (*25*) and traumatic brain injury (*24*). Our work extends this concept by showing that, in a genetically defined context, neurons can engage a selective TNF-α program that is sufficient to tune microglial activation state and synaptic engulfment. This neuron-initiated cytokine signaling is conceptually aligned with observations that TNF signaling can regulate synapse elimination and circuit stability (*41, 43, 44*), and it suggests that neuronal cytokine release should be considered not merely a secondary marker of pathology but a potential upstream instructive cue in neuron–microglia crosstalk.

UBE3A has well-established roles in synaptic plasticity and experience-dependent circuit maturation (*11, 12, 45, 46*), and its dysregulation is implicated beyond classical AS biology, including neurodevelopmental phenotypes associated with increased *UBE3A* dosage (*17, 18*) and broader immune-related contexts (*19*). A major challenge in assigning intercellular causality *in vivo* is that neuron, microglia, and astrocyte states are tightly coupled and co-evolve. Here, we leveraged a defining feature of AS, the neuron-specific imprinting of UBE3A (*13, 31, 47*), to create an experimental “separation of variables,” where a neuron-focused perturbation can be interrogated while glial UBE3A expression is relatively preserved at baseline. Within this framework, we find that neuronal UBE3A loss triggers TNF-α induction and that microglia exhibit early dysregulation in developing hippocampus alongside a conserved phagocytic/lysosomal transcriptional signature in an AS pig snRNA-seq dataset (*32*). These convergent results support a model in which neuronal UBE3A deficiency acts upstream to shape microglial inflammatory tone and engulfment programs during a critical developmental window when microglia regulate synapse refinement.

Several limitations should be acknowledged. First, our mechanistic dissection relies heavily on iPSC-derived induced neurons (iN) and microglia-like cells (iMG), which provide a tractable human platform but may not fully recapitulate region-specific maturation, developmental timing, or the multi-glial context present *in vivo* (*27, 29*). Second, Second, although neuronal UBE3A depletion was associated with increased TNFA transcripts and TNF-linked microglial inflammatory readouts, we did not directly quantify secreted TNF-α protein (e.g., by ELISA) from neuronal cultures or conditioned media; given the low basal abundance of TNF-α and the limited dynamic range of immunostaining relative to transcript assays, protein-level quantification will be important to establish physiological effect sizes. Third, because we did not perform neuronal TNF-α knockdown or neutralization, our data implicate, but do not formally prove, TNF-α as the causal neuron-derived effector; future studies should incorporate orthogonal perturbations (e.g., neuron-restricted TNF inhibition or microglia-specific TNFR manipulation) to establish source and receptor specificity within this axis. Fourth, our synapse-related readout in co-culture uses 2D colocalization of presynaptic markers with IBA1+ microglia as a proxy for microglia-associated synaptic material and cannot distinguish close apposition from true internalization; definitive assessment will require 3D compartmental analyses, lysosomal colocalization, or ultrastructural validation, consistent with established standards in the synapse elimination field. Finally, our in vivo observation of reduced microglial density alongside increased activation-associated markers raises questions about whether altered microglial survival, redistribution, or developmental maturation contributes to the net phenotype, issues that will benefit from longitudinal profiling and fate/turnover assays across developmental stages.

Our results suggest that TNF/TNFR pathway modulation can influence microglial inflammatory state and synapse-associated phenotypes downstream of neuronal UBE3A loss. TNF signaling is widely studied as a therapeutic axis across neurological diseases, including neurodegenerative and inflammatory contexts (*48, 49*). In our system, pharmacologic TNF receptor pathway inhibition (e.g., R7050) dampened microglial inflammatory outputs and normalized excessive synaptic engulfment, suggesting a potential strategy to mitigate early circuit vulnerability in UBE3A-linked neurodevelopmental disease. While AS provides a powerful model for isolating neuron-to-microglia causality, the broader implication is that neuron-initiated inflammatory programs may represent a convergent mechanism of circuit destabilization across neurodevelopmental and neuroinflammatory conditions. Further preclinical development should focus on optimizing timing, CNS target engagement, and long-term safety, with the goal of translating microglia-modulating interventions into disease-modifying treatments for neurodevelopmental disorders, led by AS.

## METHODS

### Mouse housing and strains used

All animal procedures were approved by the Brown University Institutional Animal Care and Use Committee and complied with the NIH Guide for the Care and Use of Laboratory Animals. Angelman syndrome (AS) model mice (*30*) were obtained from The Jackson Laboratory (B6.129S7–Ube3a^tm1Alb/J; JAX #016590). To generate AS mice, WT C57BL/6 mice were crossed with heterozygous B6.129S7–Ube3a^tm1Alb/J females to produce offspring with loss of maternal Ube3a expression in neurons. Genotyping was performed according to The Jackson Laboratory protocols. Mice were weaned at 3 weeks of age and housed 2–3 per cage under a 12 h light/dark cycle with ad libitum access to food and water.

### iPSC-induced neurons (iN), microglia (iMG) and co-culture

Human iPSC-derived microglia-like cells (iMG) and induced neurons (iN) were generated from the KOLF2.J1 reference iPSC line (distributed by The Jackson Laboratory) using published protocols (*29, 50, 51*). Briefly, iPSCs infected with both FUW-rtTA and pTetO-CEBPA-T2A-SPI1-T2A-Puro were selected by Puromycin and differentiated under cytokine combinations (*50, 51*).

iN generation. iNs were generated by forced expression of Neurogenin-2 (NGN2) (*27, 28, 52*). Briefly, NGN2-induced cells were selected with puromycin (2–5 μg/mL) and maintained in Neurobasal medium (Thermo Fisher Scientific #10888022) supplemented with N2 (Thermo Fisher Scientific #17502001) and B27 (Thermo Fisher Scientific #17504044).

iMG generation. iPSCs were transduced with FUW-rtTA and pTetO-CEBPA-T2A-SPI1-T2A-Puro, selected with puromycin, and differentiated using defined cytokine combinations as previously described (*29*).

siRNA knockdown in neurons. For neuronal UBE3A knockdown, siRNA targeting UBE3A (siUBE3A) or a non-targeting control (siNC) was applied to iNs at ∼DIV8 at a final concentration of 100 nM and maintained for 2–3 days prior to downstream assays.

Neuron–microglia co-culture. iNs were plated on coated 24-well plates containing coverslips for imaging and on 12-well plates for RNA analysis. iMG were added at DIV8–9 to ∼DIV15 iNs (23,31). For neuronal labeling, lentivirus expressing GFP (FUGW; Addgene #14883) was added to iNs at ∼DIV12. For co-culture experiments requiring neuronal knockdown at later stages, iNs were transfected with siNC or siUBE3A at ∼DIV14. Co-cultures were harvested at indicated time points.

### HMC3 cells

HMC3 cells (*37*) (ATCC #CRL-3304) were maintained in EMEM (ATCC #30-2003) supplemented with 10% FBS (ATCC #30-2020). Cells were passaged at a 1:5 ratio in 10 cm dishes and plated at 1 × 10^5 cells per well in 12-well plates for experiments. Treatments were initiated when cultures reached ∼70% confluence.

### Quantitative real-time PCR (qPCR)

Total RNA was extracted using the RNeasy Mini Kit (Qiagen #74104). cDNA was synthesized from 500 ng RNA using the iScript™ gDNA Clear cDNA Synthesis Kit (Bio-Rad #1725035) to remove genomic DNA contamination. qPCR was performed on a CFX96 Touch Real-Time PCR Detection System using cycling conditions of 95°C for 15 s and 60°C for 1 min for 40 cycles, followed by melt-curve analysis. Relative expression was calculated using the 2^−ΔΔCT method, normalized to GAPDH, and expressed as fold change relative to control samples. FAM probe assays for UBE3A, IBA1, C3, IL1B, TNFA, ADAM10, CD45, TUBB3, and GAPDH were purchased from Integrated DNA Technologies (IDT).

### Immunofluorescent staining

Mouse tissue. Mice were deeply anesthetized (ketamine/xylazine, 100/10 mg/kg) and perfused with ice-cold PBS followed by 4% paraformaldehyde (PFA; Millipore Sigma P6148) in PBS. Brains were post-fixed overnight in 4% PFA at 4°C, cryoprotected in 15% then 30% sucrose, embedded in OCT (VWR #25608-930), and sectioned coronally at 25 μm. Every fifth section was collected on charged slides (Globe Scientific 1354W). Sections were air-dried for ≥2 h, washed in PBS to remove OCT, and blocked for 30 min in 5% normal goat serum (Fisher Scientific 16-210-064).

Primary antibodies were diluted in 5% normal goat serum containing 0.1% Triton X-100 (Fisher Scientific PI85111) and incubated for 4 h at room temperature or overnight at 4°C: rat anti–TNF-α (R&D Systems #1AF-510-SP, 1:100), rabbit anti–Complement C3 (Thermo Fisher Scientific #PA5-21349, 1:200), rat anti-CD68 (Abcam ab53444, 1:100), chicken anti-Synapsin 1 (Aves #106009, 1:500), and chicken anti–Synaptotagmin 1/2 (Synaptic Systems #105003, 1:1000). After PBS washes, sections were incubated for 3 h at room temperature with secondary antibodies diluted in 5% normal goat serum + 0.1% Triton X-100 (e.g., Thermo Fisher Scientific A11004, A32731, A21450). Slides were mounted with Fluoromount-G with DAPI (Thermo Fisher Scientific 00-4959-52).

Cell cultures. Coverslip cultures were fixed in 4% PFA for 10 min at room temperature and stained using the same blocking/incubation framework as above.

Images were acquired using a BioTek Gen5 imaging system or a Zeiss LSM 880 confocal microscope. Fluorescence intensities were quantified in ImageJ/Fiji. For animal tissue, 3–4 images per animal were quantified and averaged to yield one value per animal.

### Immunoblotting

Cells were lysed on ice in buffer containing 1% SDS, 0.5% deoxycholate, 50 mM sodium phosphate, 150 mM NaCl, 2 mM EDTA, 50 mM NaF, 10 mM sodium pyrophosphate, 1 mM sodium orthovanadate, 1 mM PMSF, 1% IGEPAL, and protease inhibitor cocktail. Lysates were centrifuged at 10,000 × g for 5 min at 4°C and supernatants were collected. Protein concentration was determined by BCA assay and samples were mixed with 4× Laemmli buffer, boiled for 10 min, resolved by SDS–PAGE, and transferred using the Bio-Rad Trans-Blot system. GAPDH, β1-tubulin, or β-actin served as loading controls.

### KEGG analysis

Pathway enrichment. Differentially expressed genes were analyzed for Kyoto Encyclopedia of Genes and Genomes (KEGG) pathway enrichment using the clusterProfiler package (*53*). Genes were mapped to KEGG pathway annotations, and enriched pathways were identified using an adjusted significance threshold of p.adjust ≤0.01.

Pathway visualization. Significantly enriched pathways were visualized with the pathview package (*54*). For each selected pathway, gene-level expression changes (log₂ fold change) were overlaid onto the corresponding KEGG pathway diagram to generate publication-quality pathway maps.

## Statistical analysis

Graphs and statistical analyses were performed in GraphPad Prism 10. Two-group comparisons used two-tailed unpaired t tests. For ≥3 groups and/or multifactor comparisons, one-way or two-way ANOVA with post hoc multiple-comparisons testing was used unless otherwise indicated. Normality was assessed using the Shapiro–Wilk test (P > 0.05 considered normal). For immunoblots, a coefficient of variation (CV) was calculated for internal control samples; experimental values were accepted when internal control CV < 10%. All results are presented as mean ± S.E.M., n means independent experiments or animal numbers, and the significance is indicated by * *p* < 0.05, ** *p* < 0.01, *** *p* < 0.001, or\*\*\*\**P* < 0.0001.

## ETHICS DECLARATIONS

### Ethics approval and consent to participate

All mouse procedures were approved by the Brown University Institutional Animal Care and Use Committee (IACUC) and were conducted in accordance with the National Institutes of Health Guide for the Care and Use of Laboratory Animals. Human induced pluripotent stem cell (iPSC) lines used in this study were obtained from established repositories and were used in accordance with the applicable institutional oversight and material transfer agreements. No human participants were recruited and no identifiable human data were collected for this study.

### Availability of data and materials

No new sequencing datasets were generated in this study. Raw data, representative images, and analysis files are available from the corresponding author upon reasonable request. Publicly available datasets reanalyzed in this study are cited in the manuscript.

## Funding

This work was supported by grants from the Foundation for Angelman Syndrome Therapeutics (PD2023-001 to X.Y., and FT2024-001 to Y.A.H.) and from the National Institutes of Health (R01AG083943 to Y.A.H.). Y.A.H. is the endowed James and Dorothy Goodman Assistant Professor of Molecular Biology, Cell Biology and Biochemistry at Brown University.

This work is the result of NIH funding, in whole or in part, and is subject to the NIH Public Access Policy. Through acceptance of this federal funding, the NIH has been given a right to make the work publicly available in PubMed Central.

### Authors’ contributions

Conceptualization: XY, YAH; Methodology: XY, YAH; Investigation: XY, XZ, YAH; Visualization: XY, XZ, YAH; Funding acquisition: XY, YAH; Project administration: XY, YAH; Writing – original draft: XY and YAH.

### Declaration of generative AI and AI-assisted technologies in the writing process

During the preparation of this work the authors used Hemingway Editor in order to correct grammar errors and minor language issues for the first draft. After using this tool/service, the authors reviewed and edited the content as needed and take full responsibility for the content of the publication.

---

**Figure S1.**
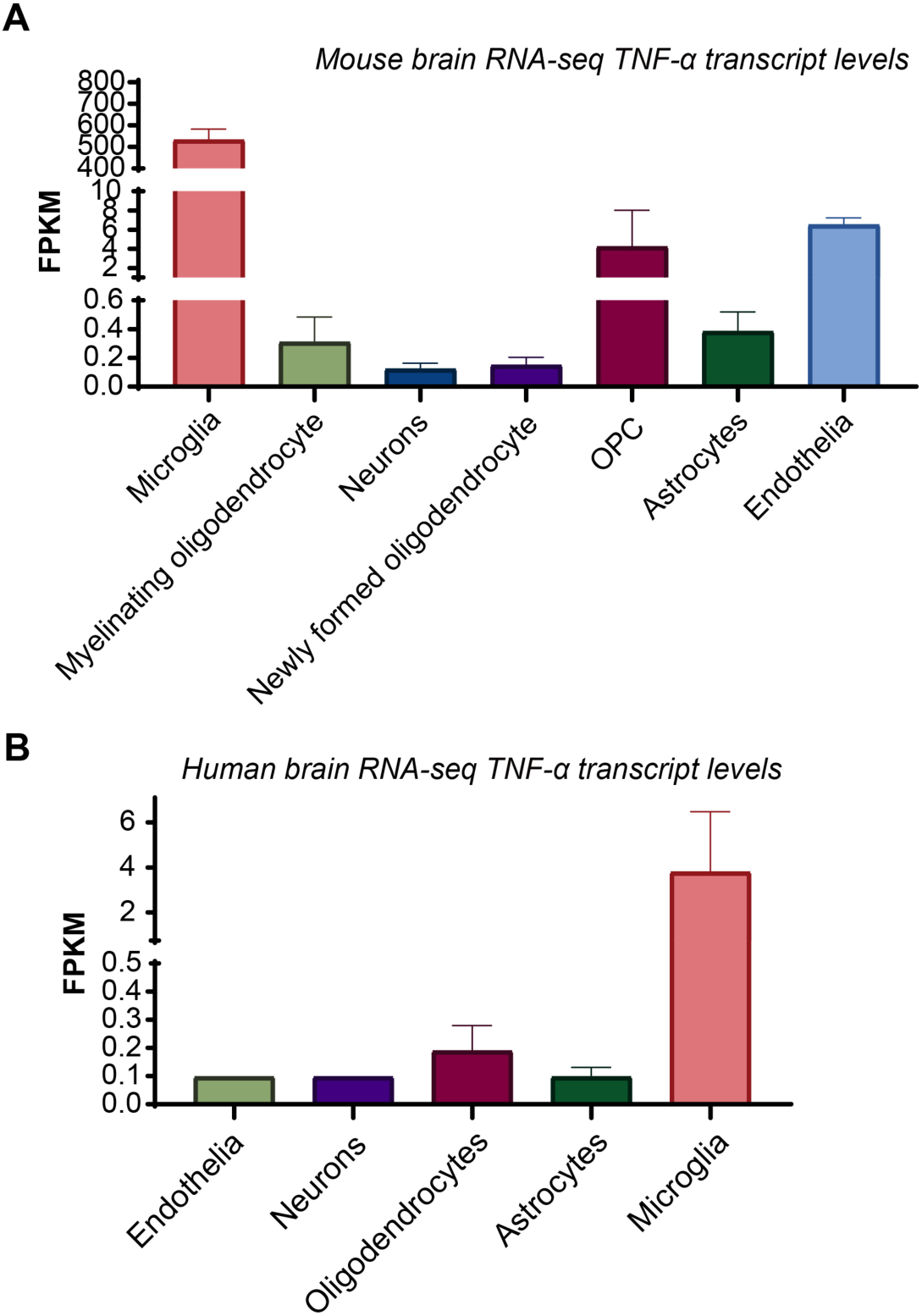
(related to Fig. 1) TNF-α is enriched in microglia but is also detectable in neurons across mouse and human brain datasets. (A) Cell type–specific *Tnf* expression in mouse brain, re-plotted from the transcriptomic dataset reported by Zhang et al. (2014). (B) Cell type–specific TNF expression in human brain, re-plotted from the transcriptomic dataset reported by Zhang et al. (2016).

**Figure S2.**
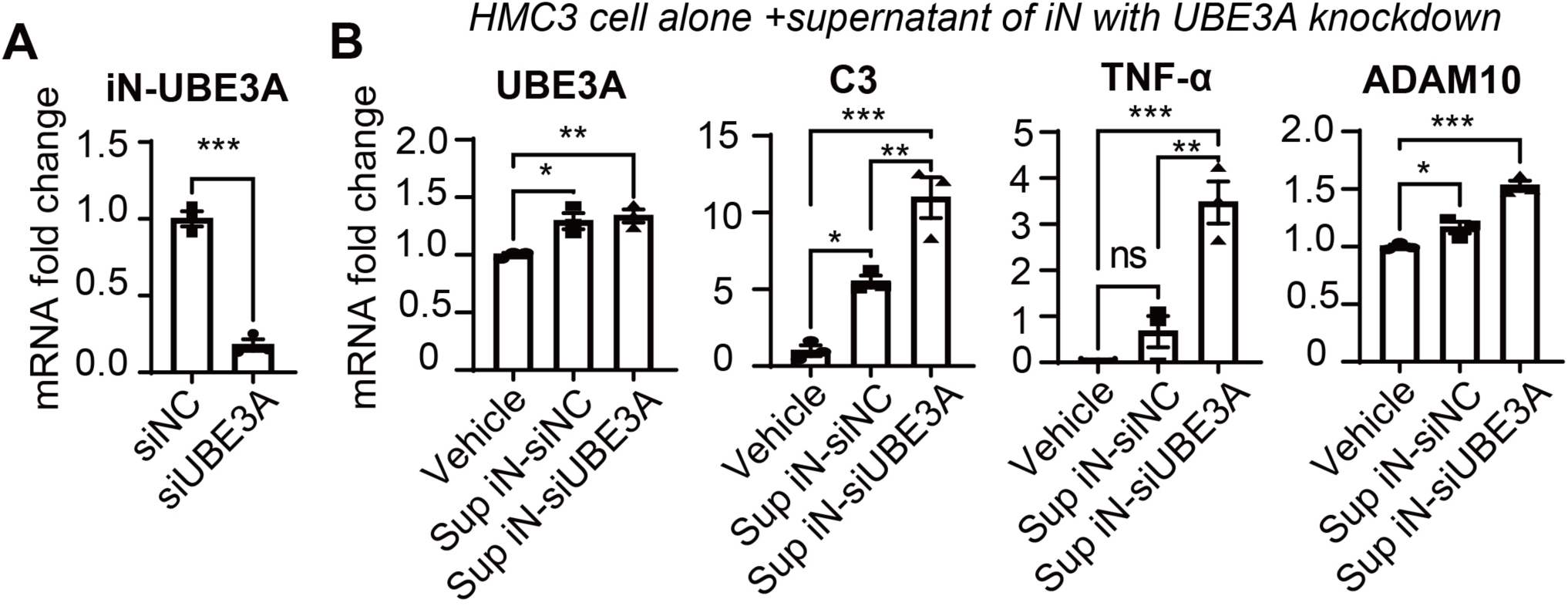
(related to Fig. 1) Neuron-conditioned media from UBE3A-deficient neurons elicits inflammatory gene induction in the HMC3 human microglial cell line. To parallel the conditioned-media paradigm in Fig. 1D using an independent microglial model, HMC3 cells were treated with supernatant collected from human induced neurons (iN) transduced with non-targeting control siRNA (siNC) or siRNA targeting UBE3A (siUBE3A). (A) qPCR validation of UBE3A knockdown in DIV14 iN, measured 2 days after siRNA transduction. Statistics: two-tailed unpaired t test. **** p < 0.0001. (B) qPCR analysis of HMC3 cells following 24-hour stimulation with neuron-conditioned media, assessing UBE3A and inflammation-associated transcripts (C3, TNFá, ADAM10). Statistics: one-way ANOVA. * p< 0.05, ** p< 0.01, *** p< 0.001, **** p< 0.0001. n = 4 independent experiments. Data are mean ± SEM

**Figure S3.**
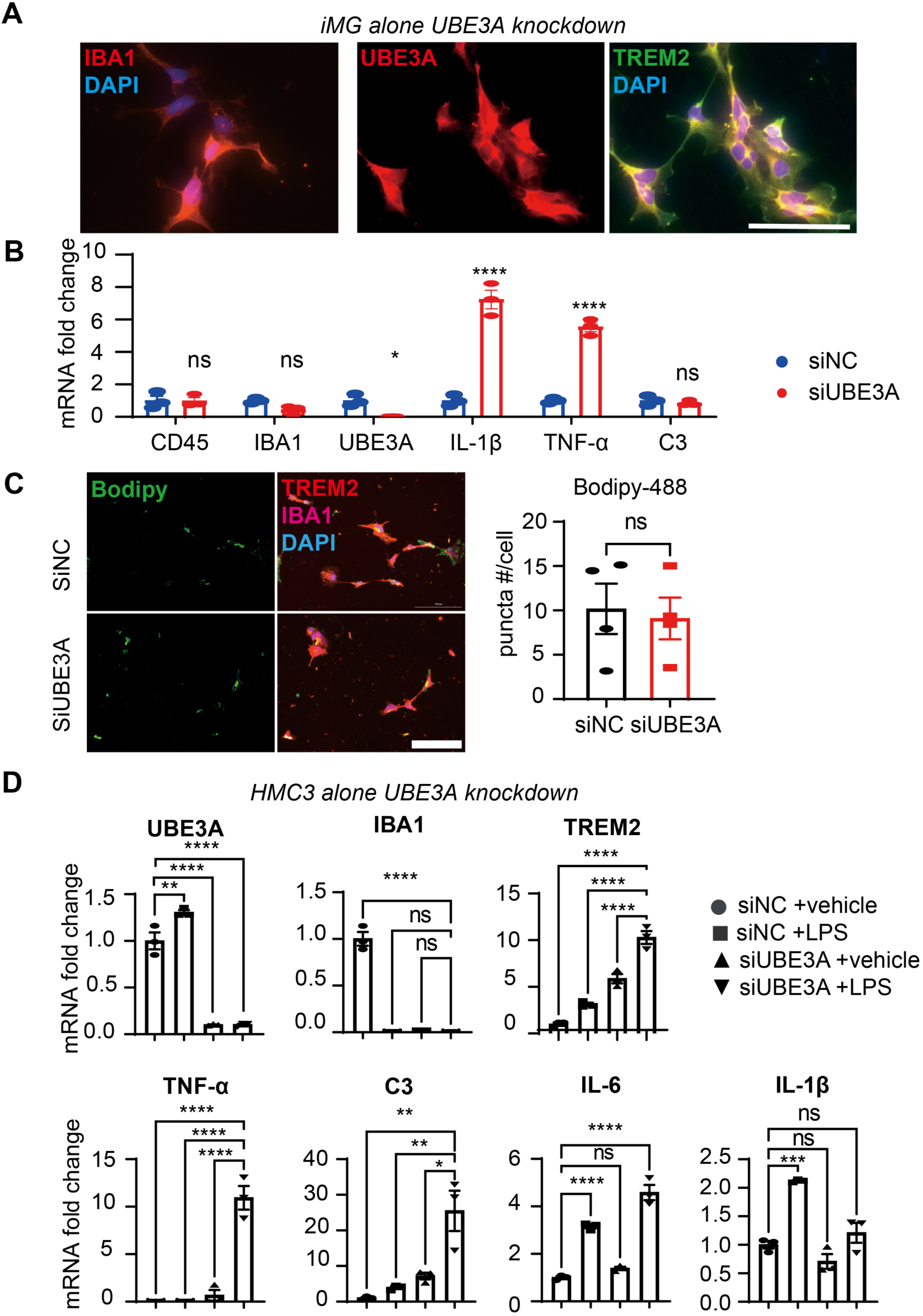
(related to Fig. 1) Cell-autonomous UBE3A knockdown increases inflammatory tone in human microglia-like cells without disrupting microglial identity or basal lipid-droplet burden. (A) Representative immunofluorescence images of iPSC-induced microglia-like cells (iMG) transduced with non-targeting control siRNA (siNC) or siRNA targeting UBE3A (siUBE3A), stained for UBE3A and microglial markers IBA1 and TREM2. Nuclei are counterstained with DAPI. (B) qPCR analysis of siNC- or siUBE3A-treated iMG showing that UBE3A knockdown elevates pro-inflammatory cytokine transcripts (IL1B and TNFA) while leaving microglial marker/identity genes (e.g., CD45, IBA1) largely unchanged. n = 3 independent cultures. Statistics: two-way ANOVA. * p< 0.05, **** p< 0.0001. (C) BODIPY 493/503 (BODIPY-488) staining and quantification of intracellular neutral lipid droplet puncta in iMG following siNC or siUBE3A treatment. This assay reports lipid droplet accumulation, a metabolic feature that can accompany microglial activation states; UBE3A knockdown did not significantly alter basal lipid-droplet burden. n = 4 independent cultures. Statistics: two-tailed unpaired t test. (D) qPCR analysis of the HMC3 human microglial cell line transduced with siNC or siUBE3A and stimulated ± LPS, measuring UBE3A and microglial markers (TREM2, IBA1) together with inflammatory transcripts (IL6, IL1B, TNFA, C3). UBE3A knockdown alone produced minimal changes at baseline but potentiated LPS-induced inflammatory gene induction, consistent with microglial priming. n = 3 independent cultures. Statistics: Owo-way ANOVA. * p< 0.05, **** p< 0.0001.

**Figure S4.**
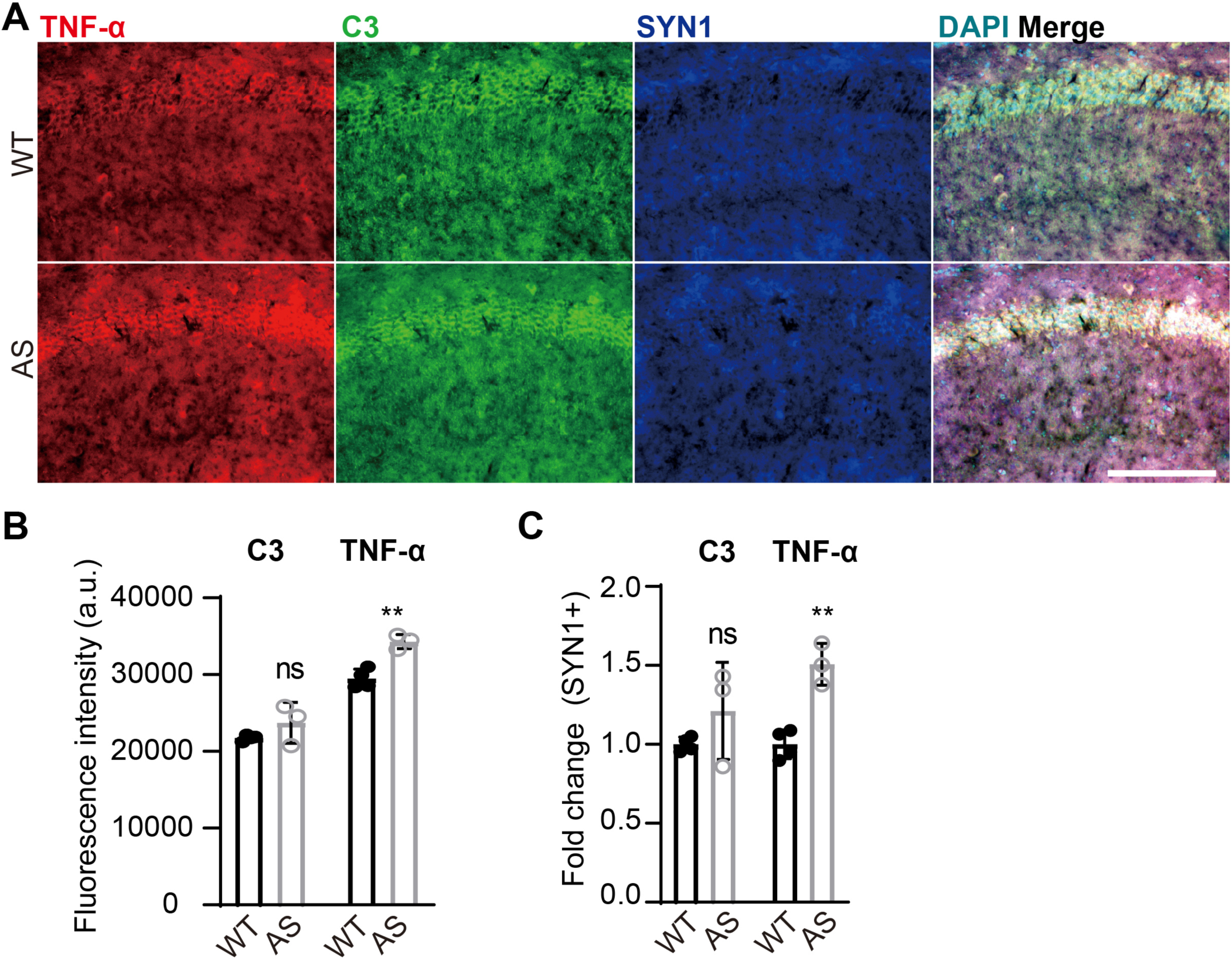
(related to Fig. 2) TNF-α and C3 are elevated in AS hippocampus, with prominent increases within SYN1⁺ neuronal regions. (A) Representative immunofluorescence images of hippocampal sections from WT and Angelman syndrome (AS) mice stained for C3 (green), the neuronal/synaptic marker SYN1 (blue), and TNF-α (red). Scale bar, 200 μm. (B) Quantification of total fluorescence intensity for C3 and TNF-α across the imaged hippocampal fields. (C) Quantification of C3 and TNF-α signal intensity restricted to SYN1⁺ regions (neuronal/synaptic areas), indicating that the relative increase in these inflammatory signals is enriched within neuronal compartments in AS. n = 4 WT and 3 AS animals. Statistics: two-way ANOVA. * p< 0.05. Data are mean ± SEM.

**Figure S5.**
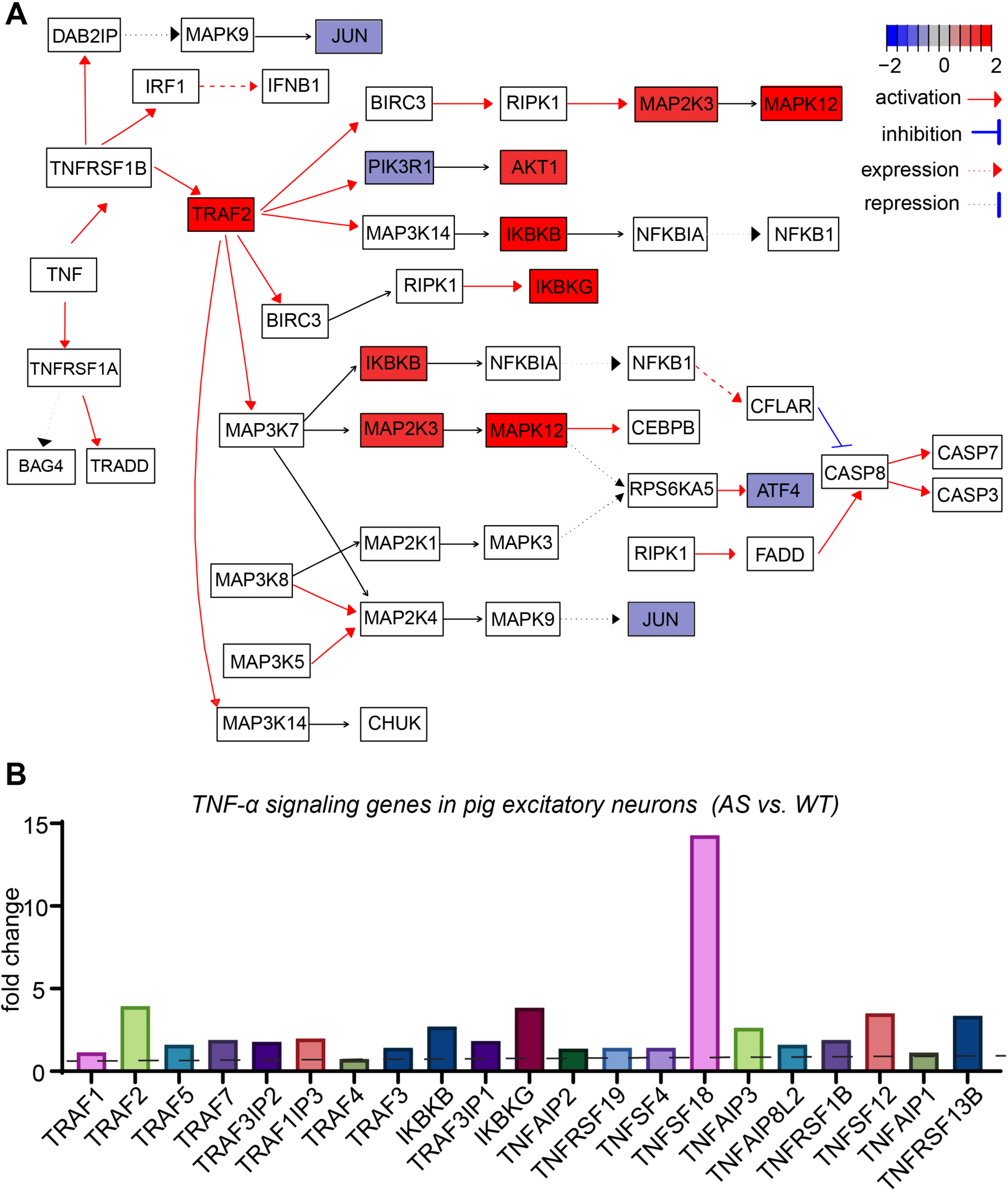
(related to Fig. 2) Reanalysis of AS pig snRNA-seq reveals TNF-α signaling pathway activation in excitatory neurons. (A) KEGG pathway enrichment analysis of differentially expressed genes (AS vs. WT) within the excitatory neuron cluster extracted from a published maternal-deletion AS pig frontal cortex single-nucleus RNA-seq dataset (Coffell et al., 2025). Enriched pathways highlight activation of TNF-α signaling in AS excitatory neurons. (B) Bar-graph representation of significantly upregulated TNF-α pathway–associated genes in AS excitatory neurons relative to WT, plotted from the same subsetted excitatory neuron dataset.

**Figure S6.**
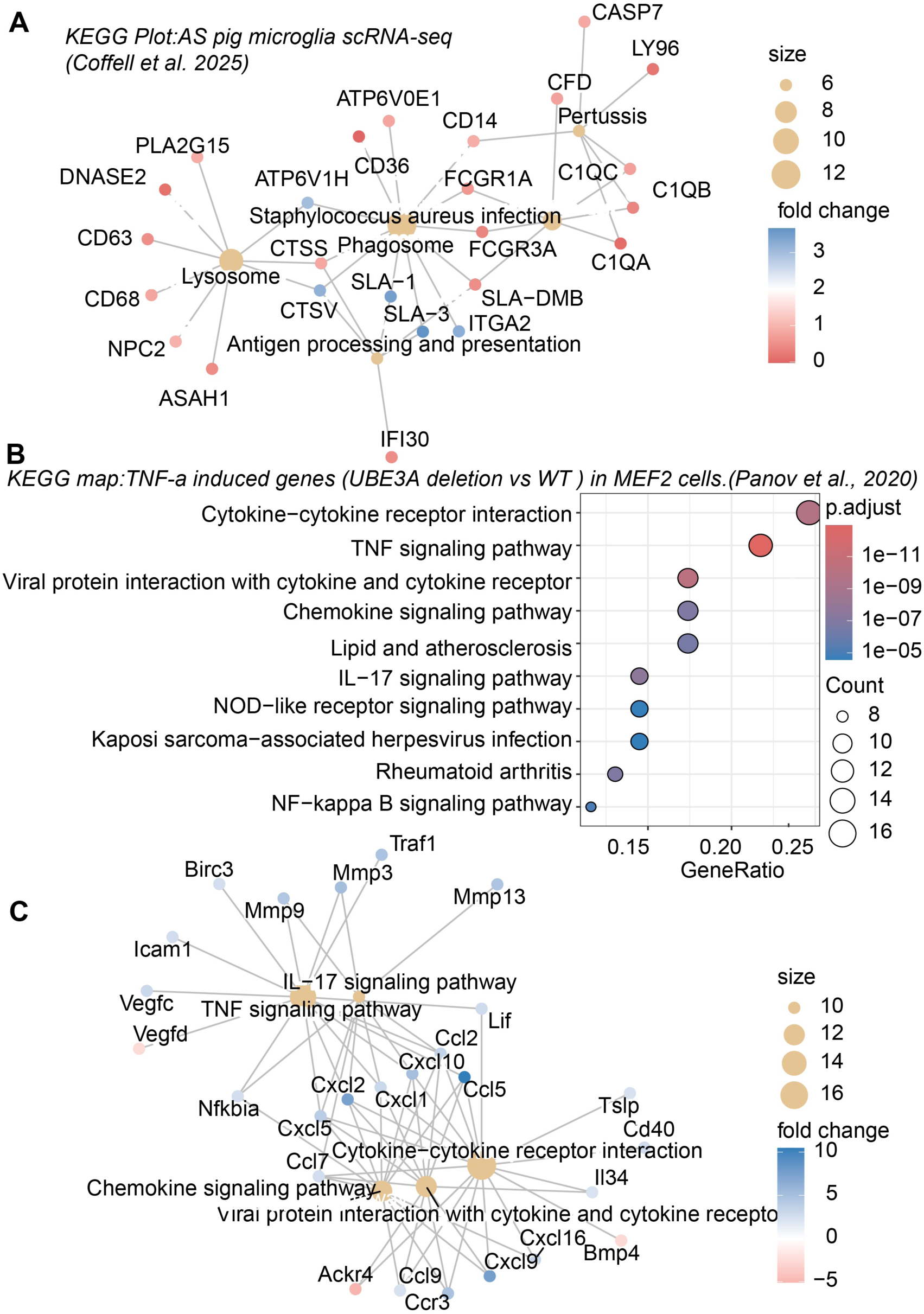
(related to Fig. 3) Transcriptome analyses link microglial phagocytic/lysosomal programs in the AS pig brain to TNF-responsive inflammatory gene networks. (A) KEGG gene–pathway network plot generated from differentially expressed genes (AS vs. WT) in microglia subsetted from a published maternal-deletion AS pig brain single-nucleus RNA-seq dataset (Coffell et al., 2025). Enriched KEGG terms highlight lysosome, phagosome, and antigen processing and presentation pathways, with representative genes including complement components (e.g., C1QA/B/C), Fc receptors (FCGR1A/FCGR3A), phagocytic/innate sensors (CD14, LY96), lysosomal/phagolysosomal markers (CD68, CD63, ATP6V0/ATP6V1 subunits, CTSS/CTSV), and antigen processing and presentation MHC (SLA, pig version). Node color indicates direction and magnitude of expression change (log2 fold change; scale shown), and node size reflects network connectivity/term–gene membership as displayed in the plot legend. (B) KEGG enrichment dot plot derived from RNA-seq of *Ube3a*−/− versus *Ube3a*+/+ mouse embryonic fibroblasts (MEFs) following TNF-α stimulation (Panov *et al.*, 2020; 25 ng/mL TNF-α for 16 h). Dots represent enriched pathways among TNF-α–responsive differentially expressed genes; dot color indicates adjusted p value (p.adjust), dot size indicates the number of genes contributing to each term (Count), and the x-axis indicates GeneRatio. (C) KEGG gene–pathway network plot for significantly altered TNF-α–responsive genes from the same MEF dataset (Panov et al., 2020), highlighting induction of cytokine/chemokine programs (e.g., CXCL1/2/10, CCL5/7) and canonical inflammatory signaling modules (TNF signaling, NF-κB signaling, IL-17 signaling, and cytokine–cytokine receptor interaction). Node color indicates log2 fold change (scale shown), and node size reflects gene–term connectivity as displayed in the plot legend.

**Figure S7.**
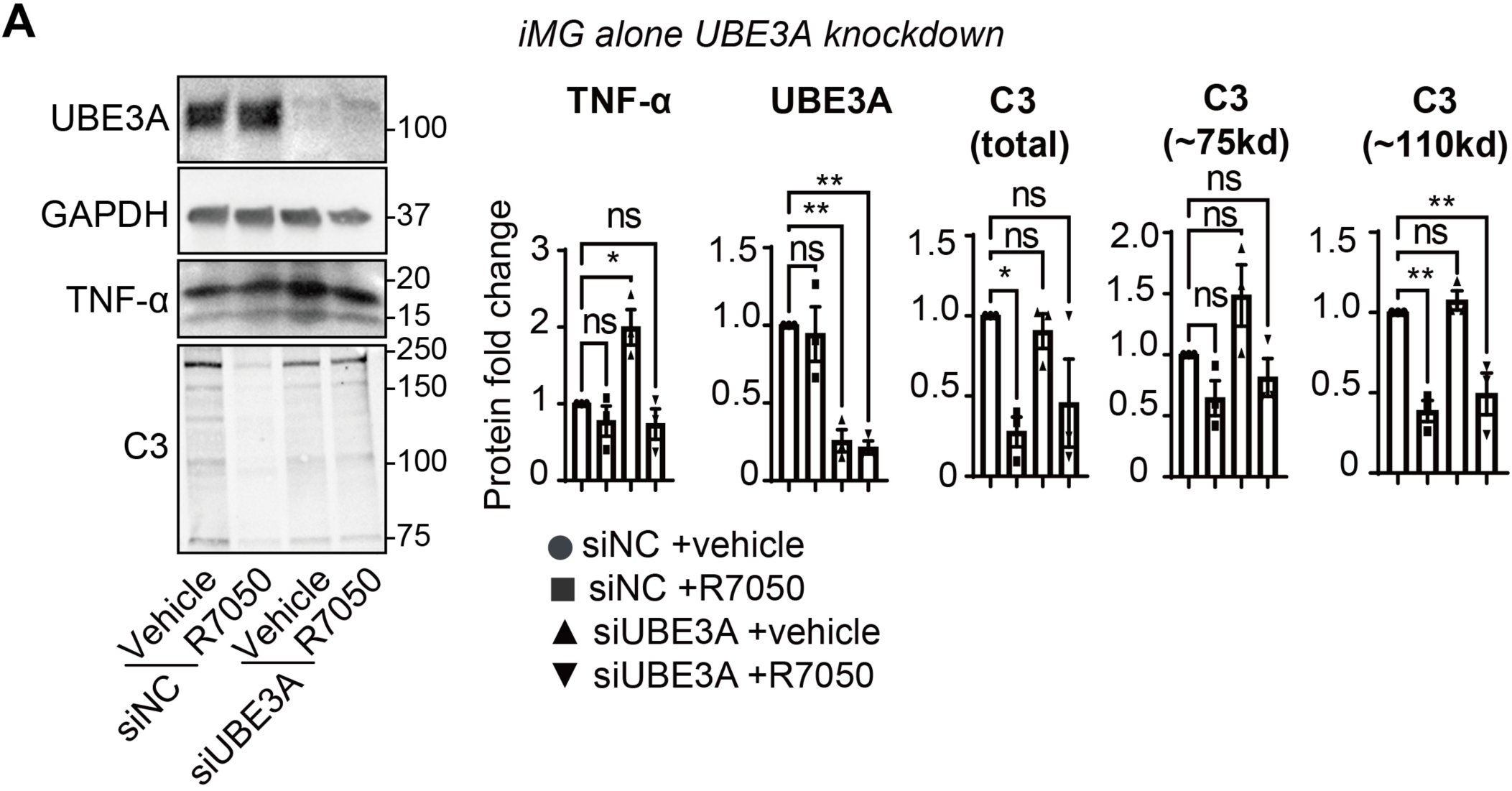
(related to Fig. 4) R7050 attenuates inflammatory protein induction in iMG with cell-autonomous UBE3A knockdown Human iMG) were transduced with non-targeting control siRNA (siNC) or siRNA targeting UBE3A (siUBE3A) and then treated with vehicle or the TNF receptor pathway inhibitor R7050. Cell lysates were analyzed by immunoblotting for UBE3A (to validate knockdown), TNF-α, and C3, with GAPDH as a loading control. Representative blots are shown on the left. Bar graphs (right) show densitometric quantification of TNF-α, UBE3A, and C3, including total C3 signal and two prominent C3 immunoreactive species (∼75 kDa and ∼110 kDa). n=4 independent experiments. *P<0.05, **P<0.01. One-way ANOVA.

## REFERENCES

1. J. R. Lukens, U. B. Eyo, Microglia and Neurodevelopmental Disorders. Annu Rev Neurosci 45, 425–445 (2022).

2. D. P. Schafer, B. Stevens, Microglia Function in Central Nervous System Development and Plasticity. Cold Spring Harb Perspect Biol 7, a020545 (2015).

3. K. Borst, A. A. Dumas, M. Prinz, Microglia: Immune and non-immune functions. Immunity 54, 2194–2208 (2021).

4. R. C. Paolicelli, G. Bolasco, F. Pagani, L. Maggi, M. Scianni, P. Panzanelli, M. Giustetto, T. A. Ferreira, E. Guiducci, L. Dumas, D. Ragozzino, C. T. Gross, Synaptic pruning by microglia is necessary for normal brain development. Science 333, 1456–1458 (2011).

5. M. W. Salter, B. Stevens, Microglia emerge as central players in brain disease. Nat Med 23, 1018–1027 (2017).

6. D. P. Schafer, E. K. Lehrman, A. G. Kautzman, R. Koyama, A. R. Mardinly, R. Yamasaki, R. M. Ransohoff, M. E. Greenberg, B. A. Barres, B. Stevens, Microglia sculpt postnatal neural circuits in an activity and complement-dependent manner. Neuron 74, 691–705 (2012).

7. C. M. Sellgren, J. Gracias, B. Watmuff, J. D. Biag, J. M. Thanos, P. B. Whittredge, T. Fu, K. Worringer, H. E. Brown, J. Wang, A. Kaykas, R. Karmacharya, C. P. Goold, S. D. Sheridan, R. H. Perlis, Increased synapse elimination by microglia in schizophrenia patient-derived models of synaptic pruning. Nat Neurosci 22, 374–385 (2019).

8. J. Wu, J. Zhang, X. Chen, K. Wettschurack, Z. Que, B. A. Deming, M. I. Olivero-Acosta, N. Cui, M. Eaton, Y. Zhao, S. M. Li, M. Suzuki, I. Chen, T. Xiao, M. S. Halurkar, P. Mandal, C. Yuan, R. Xu, W. A. Koss, D. Du, F. Chen, L. J. Wu, Y. Yang, Microglial over-pruning of synapses during development in autism-associated SCN2A-deficient mice and human cerebral organoids. Mol Psychiatry 29, 2424–2437 (2024).

9. G. Tang, K. Gudsnuk, S. H. Kuo, M. L. Cotrina, G. Rosoklija, A. Sosunov, M. S. Sonders, E. Kanter, C. Castagna, A. Yamamoto, Z. Yue, O. Arancio, B. S. Peterson, F. Champagne, A. J. Dwork, J. Goldman, D. Sulzer, Loss of mTOR-dependent macroautophagy causes autistic-like synaptic pruning deficits. Neuron 83, 1131–1143 (2014).

10. H. J. Kim, M. H. Cho, W. H. Shim, J. K. Kim, E. Y. Jeon, D. H. Kim, S. Y. Yoon, Deficient autophagy in microglia impairs synaptic pruning and causes social behavioral defects. Mol Psychiatry 22, 1576–1584 (2017).

11. X. Yang, Characterizing spine issues: If offers novel therapeutics to Angelman syndrome. Dev Neurobiol 80, 200–209 (2020).

12. K. Yashiro, T. T. Riday, K. H. Condon, A. C. Roberts, D. R. Bernardo, R. Prakash, R. J. Weinberg, M. D. Ehlers, B. D. Philpot, Ube3a is required for experience-dependent maturation of the neocortex. Nat Neurosci 12, 777–783 (2009).

13. A. M. Mabb, M. C. Judson, M. J. Zylka, B. D. Philpot, Angelman syndrome: insights into genomic imprinting and neurodevelopmental phenotypes. Trends Neurosci 34, 293–303 (2011).

14. P. L. Greer, R. Hanayama, B. L. Bloodgood, A. R. Mardinly, D. M. Lipton, S. W. Flavell, T. K. Kim, E. C. Griffith, Z. Waldon, R. Maehr, H. L. Ploegh, S. Chowdhury, P. F. Worley, J. Steen, M. E. Greenberg, The Angelman Syndrome protein Ube3A regulates synapse development by ubiquitinating arc. Cell 140, 704–716 (2010).

15. H. Kim, P. A. Kunz, R. Mooney, B. D. Philpot, S. L. Smith, Maternal Loss of Ube3a Impairs Experience-Driven Dendritic Spine Maintenance in the Developing Visual Cortex. J Neurosci 36, 4888–4894 (2016).

16. J. Sun, G. Zhu, Y. Liu, S. Standley, A. Ji, R. Tunuguntla, Y. Wang, C. Claus, Y. Luo, M. Baudry, X. Bi, UBE3A Regulates Synaptic Plasticity and Learning and Memory by Controlling SK2 Channel Endocytosis. Cell Rep 12, 449–461 (2015).

17. A. Noor, L. Dupuis, K. Mittal, A. C. Lionel, C. R. Marshall, S. W. Scherer, T. Stockley, J. B. Vincent, R. Mendoza-Londono, D. J. Stavropoulos, 15q11.2 Duplication Encompassing Only the UBE3A Gene Is Associated with Developmental Delay and Neuropsychiatric Phenotypes. Hum Mutat 36, 689–693 (2015).

18. B. Roy, E. Amemasor, S. Hussain, K. Castro, UBE3A: The Role in Autism Spectrum Disorders (ASDs) and a Potential Candidate for Biomarker Studies and Designing Therapeutic Strategies. Diseases 12, (2023).

19. X. Yang, Y. A. Huang, Unraveling the Roles of UBE3A in Neurodevelopment and Neurodegeneration. Int J Mol Sci 26, (2025).

20. S. A. Liddelow, K. A. Guttenplan, L. E. Clarke, F. C. Bennett, C. J. Bohlen, L. Schirmer, M. L. Bennett, A. E. Münch, W. S. Chung, T. C. Peterson, D. K. Wilton, A. Frouin, B. A. Napier, N. Panicker, M. Kumar, M. S. Buckwalter, D. H. Rowitch, V. L. Dawson, T. M. Dawson, B. Stevens, B. A. Barres, Neurotoxic reactive astrocytes are induced by activated microglia. Nature 541, 481–487 (2017).

21. M. Linnerbauer, M. A. Wheeler, F. J. Quintana, Astrocyte Crosstalk in CNS Inflammation. Neuron 108, 608–622 (2020).

22. G. Olmos, J. Lladó, Tumor necrosis factor alpha: a link between neuroinflammation and excitotoxicity. Mediators Inflamm 2014, 861231 (2014).

23. T. Balzano, Y. M. Arenas, S. Dadsetan, J. Forteza, S. Gil-Perotin, L. Cubas-Nunez, B. Casanova, F. Gracia, N. Varela-Andres, C. Montoliu, M. Llansola, V. Felipo, Sustained hyperammonemia induces TNF-a IN Purkinje neurons by activating the TNFR1-NF-kappaB pathway. J Neuroinflammation 17, 70 (2020).

24. S. M. Knoblach, L. Fan, A. I. Faden, Early neuronal expression of tumor necrosis factor-alpha after experimental brain injury contributes to neurological impairment. J Neuroimmunol 95, 115–125 (1999).

25. T. Liu, R. K. Clark, P. C. McDonnell, P. R. Young, R. F. White, F. C. Barone, G. Z. Feuerstein, Tumor necrosis factor-alpha expression in ischemic neurons. Stroke 25, 1481–1488 (1994).

26. J. L. Tchelingerian, J. Quinonero, J. Booss, C. Jacque, Localization of TNF alpha and IL-1 alpha immunoreactivities in striatal neurons after surgical injury to the hippocampus. Neuron 10, 213–224 (1993).

27. K. Connolly, M. Lehoux, B. Assetta, Y. A. Huang, Modeling Cellular Crosstalk of Neuroinflammation Axis by Tri-cultures of iPSC-Derived Human Microglia, Astrocytes, and Neurons. Methods Mol Biol 2683, 79–87 (2023).

28. Y. A. Huang, B. Zhou, M. Wernig, T. C. Sudhof, ApoE2, ApoE3, and ApoE4 Differentially Stimulate APP Transcription and Abeta Secretion. Cell 168, 427-441.e421 (2017).

29. M. Lehoux, K. Connolly, B. Assetta, Y. A. Huang, The Generation and Functional Characterization of Human Microglia-Like Cells Derived from iPS and Embryonic Stem Cells. Methods Mol Biol 2683, 69–78 (2023).

30. Y. H. Jiang, D. Armstrong, U. Albrecht, C. M. Atkins, J. L. Noebels, G. Eichele, J. D. Sweatt, A. L. Beaudet, Mutation of the Angelman ubiquitin ligase in mice causes increased cytoplasmic p53 and deficits of contextual learning and long-term potentiation. Neuron 21, 799–811 (1998).

31. X. Yang, Towards an understanding of Angelman syndrome in mice studies. J Neurosci Res 98, 1162–1173 (2020).

32. A. Coffell, L. Schuller, S. Christian, S. V. Dindot, Cell-type-specific dysregulated gene expression in the frontal cortex of an Angelman syndrome pig model. bioRxiv, 2025.2008.2020.671338 (2025).

33. L. S. Myers, S. G. Christian, S. Simpson, R. Sper, C. Taylor, L. Montes, T. B. C. Jepp, D. Ramos, L. Schuller, K. Konganti, W. Friedeck, O. Habib, M. Hodge, A. J. Taylor, A. Coffell, A. Schlafer, M. Matt, B. Revell, C. Knight, C. C. Barreña, W. J. Murphy, E. J. Weeber, D. J. Segal, A. Anderson, K. R. Nash, J. L. Silverman, J. A. Piedrahita, S. V. Dindot, A preclinical pig model of Angelman syndrome mirrors the early developmental trajectory of the human condition. Proc Natl Acad Sci U S A 122, e2505152122 (2025).

34. D. M. Ramos, W. C. Skarnes, A. B. Singleton, M. R. Cookson, M. E. Ward, Tackling neurodegenerative diseases with genomic engineering: A new stem cell initiative from the NIH. Neuron 109, 1080–1083 (2021).

35. Y. Zhang, K. Chen, S. A. Sloan, M. L. Bennett, A. R. Scholze, S. O’Keeffe, H. P. Phatnani, P. Guarnieri, C. Caneda, N. Ruderisch, S. Deng, S. A. Liddelow, C. Zhang, R. Daneman, T. Maniatis, B. A. Barres, J. Q. Wu, An RNA-sequencing transcriptome and splicing database of glia, neurons, and vascular cells of the cerebral cortex. J Neurosci 34, 11929–11947 (2014).

36. Y. Zhang, S. A. Sloan, L. E. Clarke, C. Caneda, C. A. Plaza, P. D. Blumenthal, H. Vogel, G. K. Steinberg, M. S. Edwards, G. Li, J. A. Duncan, 3rd, S. H. Cheshier, L. M. Shuer, E. F. Chang, G. A. Grant, M. G. Gephart, B. A. Barres, Purification and Characterization of Progenitor and Mature Human Astrocytes Reveals Transcriptional and Functional Differences with Mouse. Neuron 89, 37–53 (2016).

37. C. Dello Russo, N. Cappoli, I. Coletta, D. Mezzogori, F. Paciello, G. Pozzoli, P. Navarra, A. Battaglia, The human microglial HMC3 cell line: where do we stand? A systematic literature review. J Neuroinflammation 15, 259 (2018).

38. J. Lier, W. J. Streit, I. Bechmann, Beyond Activation: Characterizing Microglial Functional Phenotypes. Cells 10, (2021).

39. J. Panov, L. Simchi, Y. Feuermann, H. Kaphzan, Bioinformatics Analyses of the Transcriptome Reveal Ube3a-Dependent Effects on Mitochondrial-Related Pathways. Int J Mol Sci 21, (2020).

40. M. D. King, C. H. Alleyne, Jr., K. M. Dhandapani, TNF-alpha receptor antagonist, R-7050, improves neurological outcomes following intracerebral hemorrhage in mice. Neurosci Lett 542, 92–96 (2013).

41. E. C. Beattie, D. Stellwagen, W. Morishita, J. C. Bresnahan, B. K. Ha, M. Von Zastrow, M. S. Beattie, R. C. Malenka, Control of synaptic strength by glial TNFalpha. Science 295, 2282–2285 (2002).

42. S. Habbas, M. Santello, D. Becker, H. Stubbe, G. Zappia, N. Liaudet, F. R. Klaus, G. Kollias, A. Fontana, C. R. Pryce, T. Suter, A. Volterra, Neuroinflammatory TNFalpha Impairs Memory via Astrocyte Signaling. Cell 163, 1730–1741 (2015).

43. D. Stellwagen, R. C. Malenka, Synaptic scaling mediated by glial TNF-alpha. Nature 440, 1054–1059 (2006).

44. X. Q. Fu, J. Peng, A. H. Wang, Z. G. Luo, Tumor necrosis factor alpha mediates neuromuscular synapse elimination. Cell Discov 6, 9 (2020).

45. X. Yang, Y. A. Huang, J. Marshall, UBE3A stabilization of beta-catenin preserves synaptic proteins essential for motor and cognitive functions in Angelman Syndrome. Mol Autism 16, 60 (2025).

46. X. Yang, Y. A. Huang, J. Marshall, Targeting TrkB-PSD-95 coupling to mitigate neurological disorders. Neural Regen Res 20, 715–724 (2025).

47. K. Buiting, C. Williams, B. Horsthemke, Angelman syndrome - insights into a rare neurogenetic disorder. Nat Rev Neurol 12, 584–593 (2016).

48. W. Chadwick, T. Magnus, B. Martin, A. Keselman, M. P. Mattson, S. Maudsley, Targeting TNF-alpha receptors for neurotherapeutics. Trends Neurosci 31, 504–511 (2008).

49. N. Torres-Acosta, J. H. O’Keefe, E. L. O’Keefe, R. Isaacson, G. Small, Therapeutic Potential of TNF-alpha Inhibition for Alzheimer’s Disease Prevention. J Alzheimers Dis 78, 619–626 (2020).

50. S. W. Chen, Y. H. Wong, Directed Differentiation of Human iPSCs into Microglia-Like Cells Using Defined Transcription Factors. Methods Mol Biol 2683, 53–68 (2023).

51. S. W. Chen, Y. S. Hung, J. L. Fuh, N. J. Chen, Y. S. Chu, S. C. Chen, M. J. Fann, Y. H. Wong, Efficient conversion of human induced pluripotent stem cells into microglia by defined transcription factors. Stem Cell Reports 16, 1363–1380 (2021).

52. Y. A. Huang, B. Zhou, A. M. Nabet, M. Wernig, T. C. Sudhof, Differential Signaling Mediated by ApoE2, ApoE3, and ApoE4 in Human Neurons Parallels Alzheimer’s Disease Risk. J Neurosci, (2019).

53. S. Xu, E. Hu, Y. Cai, Z. Xie, X. Luo, L. Zhan, W. Tang, Q. Wang, B. Liu, R. Wang, W. Xie, T. Wu, L. Xie, G. Yu, Using clusterProfiler to characterize multiomics data. Nature Protocols 19, 3292–3320 (2024).

54. W. Luo, C. Brouwer, Pathview: an R/Bioconductor package for pathway-based data integration and visualization. Bioinformatics 29, 1830–1831 (2013).

